# Information theory tests critical predictions of plant defense theory for specialized metabolism

**DOI:** 10.1101/2020.01.07.897389

**Authors:** Dapeng Li, Rayko Halitschke, Ian T. Baldwin, Emmanuel Gaquerel

## Abstract

Different plant defense theories have provided important theoretical guidance in explaining patterns in plant specialized metabolism, but their critical predictions remain to be tested. Here, we systematically explored the metabolomes of *Nicotiana attenuata*, from single plants to populations, as well as of closely-related species, using unbiased MS/MS analyses and processed the abundances of compound-spectrum-based MS features within an information theory framework to test critical predictions of Optimal Defense (OD) and Moving Target (MT) theories. Information components of herbivory-elicited plant metabolomes were fully consistent with the OD theory predictions and contradicted the main prediction of the MT theory. From micro- to macro-evolutionary scales, jasmonate signaling was identified as the master determinant of OD while ethylene signaling provided fine-tuning for herbivore-specific responses annotated via MS/MS molecular networks.

**One-sentence summary:** Information theory tests defense theory predictions by providing a common currency for comparison of specialized metabolomes

## Introduction

Structurally diverse specialized metabolites are central players in plants’ adaptations to their environments and in particular in their defense against enemies (*1*). The spectacular diversification of specialized metabolism found in plants inspired several decades of intense research about its multifaceted ecological functions and nucleated a long list of plant defense theories that provided important guidance to empirical studies of the evolution and ecology of plant-insect interactions (*2*). However, these plant defense theories did not follow the canonical path of hypothetical-deductive reasoning in which critical predictions are posed at the same levels of analysis (*3*) and tested experimentally to advance the next cycle of theory development (*4*).

Technical constraints limited data acquisition to specific metabolic classes and precluded the comprehensive profiling of specialized metabolites and consequently, prevented the among-taxa comparisons which were essential for theory advancement (*5*). This lack of comprehensive metabolomics data and the processing workflows to compare the metabolic space among different plant taxa in a common currency thwarted the scientific maturation of the field.

Recent advances in the field of tandem mass spectrometry (MS/MS) metabolomics are allowing for comprehensive and high-throughput characterizations of metabolic variations within and among species that comprise complete phylogenetic clades particularly when computational methods are employed for the calculation of structural similarity among complex mixtures without *a priori* chemical knowledge (*5*). This combination of analytical and computational advances provides the missing framework required for the rigorous testing of many predictions made by the long-standing ecological and evolutionary theories on metabolic diversity.

Information theory, first introduced by Claude Shannon in his seminal article in 1948, laid the groundwork for a mathematical analysis of information (*6*) which has been adopted by many fields beyond its original applications (*7, 8*). In genomics, information theory has been successfully applied to quantify sequence conservation information (*9*) and in a transcriptomics study, information theory parsed the global changes that occur in the transcriptome (*10*). In a previous study, we applied the information theory statistical framework to metabolomics to describe tissue-level metabolic specialization in plants (*11*). Here we combine an MS/MS-based workflow with the statistical framework of information theory to characterize metabolic diversity in a common currency so as to compare critical predictions of plant defense theories.

Plant defense theoretical frameworks are frequently mutually inclusive and can be classified into two groups: those that attempt to explain the distribution of plant specialized metabolites based on defensive function, such as the Optimal Defense (OD) (*12–14*), Moving Target (MT) (*15*) and Apparency (*16*) theories, whereas others seek mechanistic explanations in how variation in resource availability influences plant growth and specialized metabolite accumulations, for instance, the Carbon:Nutrient Balance (CNB) hypothesis (*17*), the Growth Rate (GR) hypothesis (*18*) and the Growth-Differentiation Balance (GDB) hypothesis (*19*). These two groups of theories are posed at different levels of analysis (*4*). However, two theories, both addressing defensive functions at the functional level have dominated the dialogue about plant constitutive and inducible defenses: the OD theory which hypothesized that plants invest their costly chemical defense only when needed, for instance, when attacked by herbivores, thus directionally allocating compounds with defensive function according to the probability of future attack (*12–14*), and, the MT hypothesis which proposes that axes of directional metabolite changes do not exist but rather that metabolites change randomly thereby creating a metabolic “moving target” which could thwart the performance of attacking herbivores. In other words, the two theories present contrasting predictions about the metabolic reconfigurations that occur after herbivore attack: the unidirectional accumulations of metabolites with defense functions (OD) versus non-directional metabolic changes (MT) (*15*). Thus, while both hypotheses acknowledge the defensive function of specialized metabolites, which may or may not be costly to produce, they differ in their predictions regarding the directionality of induced metabolic changes. The predictions of the OD theory have received the most experimental attention to date. These tests include investigations of the variations in specific compound classes with either direct or indirect defense functions across different tissues and ontogenetic stages in plants grown under both glasshouse and natural conditions (*20–26*). However, to date, the central distinguishing prediction of the two theories, namely the directionality of the metabolic changes, remains to be tested due to the lack of workflows and statistical frameworks to conduct a globally comprehensive analysis of metabolic diversity of any organism. Here we provide such an analysis.

One of the most striking characteristics of plant specialized metabolites is their extreme structural diversity at all levels ranging from single plants, populations to congeneric species (*27*). Much of the quantitative variations in specialized metabolites can be observed at a population scale whereas strong qualitative differences are commonly maintained at species levels (*28*). Plant metabolic diversity is hence a primary dimension of functional diversity reflecting adaptations to different ecological niches, in particular those that differ in the probability of attack from a broad range of insects, including specialist and generalist herbivores (*29*). Interactions with various insects are important selective pressures which since Fraenkel’s seminal article on the *raison d’être* of plant specialized metabolites (*30*), are thought to have sculpted plant metabolic pathways during evolution (*31*). Interspecies variations in specialized metabolite diversity may additionally reflect physiological tradeoffs associated with constitutive and induced plant defense strategies against herbivory, as the two are frequently negatively correlated with each other across species (*32, 33*). While being well-defended at all times may be advantageous, timely defense-related metabolic changes provide clear benefits in allowing plants to allocate valuable resources to other physiological investments (*32, 34*) and avoid collateral damage to mutualists (*35*). Additionally, these insect herbivory-induced reconfigurations of specialized metabolite production can lead to disruptive distributions in populations (*36*) and may reflect direct readouts of substantial natural variation in jasmonate (JA) signaling, with high and low JA signaling being likely maintained in a population by tradeoffs between defense to herbivores and competition with conspecies (*37*). Furthermore, specialized metabolite biosynthetic pathways can undergo rapid loss and gain transitions during evolution, resulting in patchy metabolic distribution among closely-related species (*38*). This polymorphism can be rapidly established in response to changing herbivore regimes (*39*) implying that fluctuations in herbivore communities are key factors driving metabolic heterogeneity.

Here we specifically addressed the following questions. (*i*) How are plant metabolomes reconfigured by insect herbivory? (*ii*) What principle information components of the metabolic plasticity can be quantified to test predictions of long-standing defense theories? (*iii*) Is the reprogramming of plant metabolomes conducted in an attacker-specific fashion, and if so, what roles do phytohormones play in tailoring the specific metabolic responses, and which metabolites contribute to the species-specificity of elicited defense? (*iv*) As many defense theories make predictions that scale across levels of biological organization, we asked how consistently do the elicited metabolic responses scale from intra- to inter-specific comparisons? To this end, we systematically explored the leaf metabolome of *N. attenuata*, an ecological model plant with rich specialized metabolism and its response to attack from two native herbivores, the larvae of *S. littoralis* (Sl), a generalist, and *M. sexta* (Ms), a specialist moth species. We parsed MS/MS metabolomics spectra and extracted information theory statistical descriptors to contrast predictions of OD and MT theories. Specificity maps were created to reveal the identity of key metabolites. The analysis was extended to *N. attenuata* native populations and closely-related *Nicotiana* species to further dissect co-variations between phytohormone signaling and OD inductions.

## Results

### Statistical descriptors of metabolic plasticity and information content from large-scale MS/MS data

To capture a holistic picture of the plasticity and structure on the herbivory-induced leaf metabolome of *N. attenuata*, we used a previously developed analytical and computational workflow to comprehensively collect and deconvolute high-resolution data-independent MS/MS spectra from plant extracts (*11*). This indiscriminant approach (termed MS/MS) allows constructing non-redundant compound spectra that are subsequently used for all compound-level analyses presented here. These deconvoluted plant metabolomes are highly diverse and composed of hundreds to thousands of metabolites (here, ∼500 to 1000s MS/MSs). Principle component analysis (PCA) that transforms multi-variable dataset into a reduced dimensionality space such that the main trends in the data are interpretable is frequently adopted as an exploratory technique to parse datasets such as deconvoluted metabolomes. However, dimensionality reduction loses part of the information content in the dataset, and most importantly, PCA analysis provide no quantitative information about traits that are particularly germane for ecological theory, such as: How does herbivory reconfigure diversity (eg., richness, distribution and abundance) in specialized metabolites? Which metabolites are predictors for a given herbivory-induced state? Here we consider metabolic plasticity in an information theory framework, and quantify the diversity and specialization of a metabolome based on the Shannon entropy of the metabolic frequency distribution. Using previously implemented formulae (*10*), we calculated a set of indices that allow for the quantification of metabolome diversity (Hj index), metabolic profile specialization (δj index) and metabolic specificity of individual metabolites (Si index). Additionally, we applied a relative distance plasticity index (RDPI index) to quantify metabolome inducibility by herbivory (*40*) (Fig. 1A). Within this statistical framework, we consider MS/MS spectra as basic information units and process MS/MSs relative abundances into frequency profiles from which metabolome diversity is estimated using Shannon entropy.

**Fig. 1.**
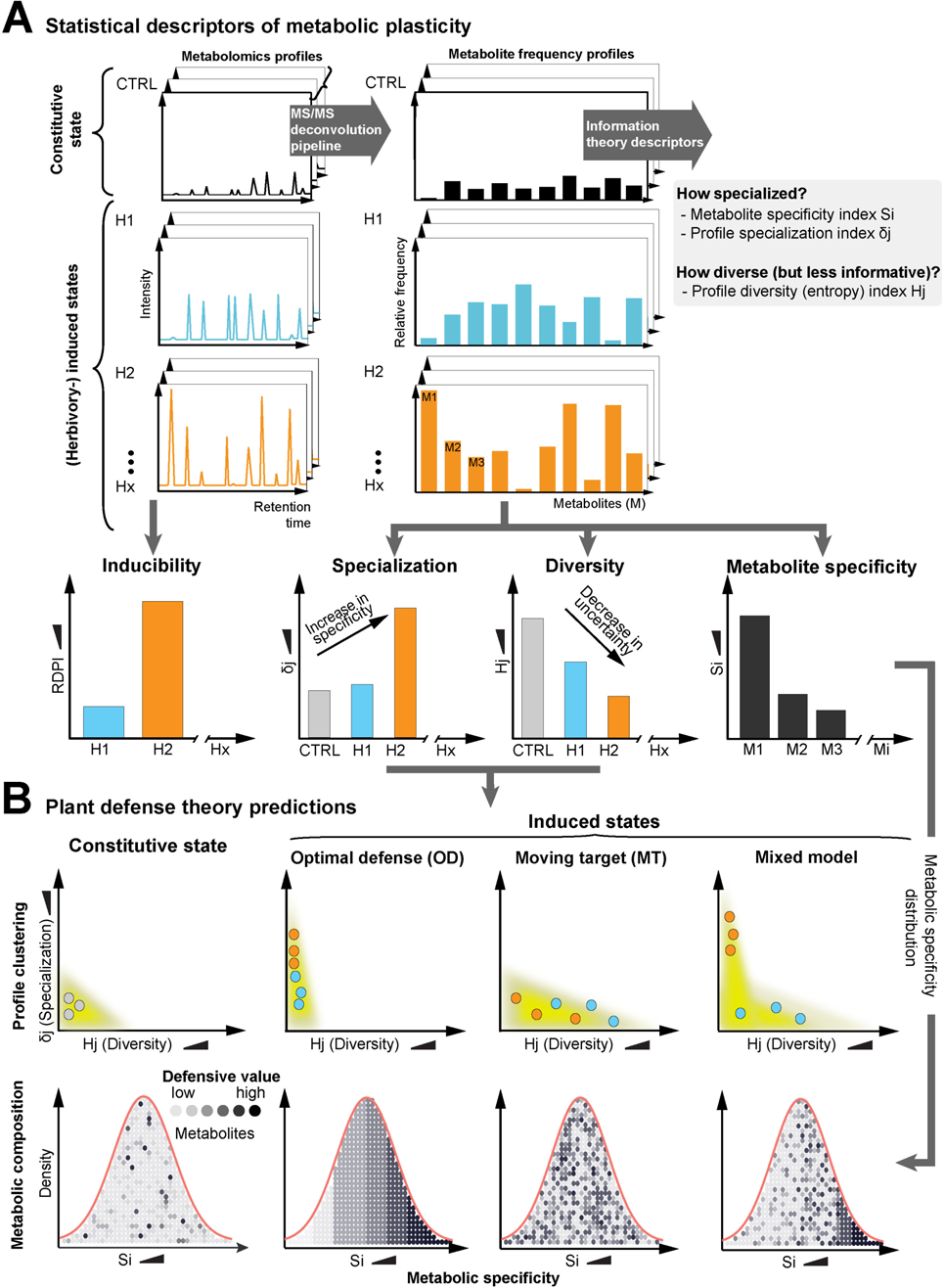
Testing and contrasting the predictions of Optimal Defense (OD) and Moving Target (MT) theories using an information theory statistical framework. (A) Scheme illustrating the extraction of statistical descriptors used to quantify metabolic plasticity, namely metabolite inducibility (relative distance plasticity index: RDPI) and three information theory descriptors—diversity (Hj index), specialization (δj index) and metabolic specificity (Si index), which were obtained by processing relative abundances of compound spectrum-based MS features derived from a large-scale collection of MS/MS features from herbivory-elicited (indicated as H_1_ to H_x_) plant extracts. An increase in specialization (δj) indicates that, on average, more herbivory-specific metabolites are produced, whereas a decrease in diversity (Hj) indicates that either fewer metabolites are produced or that the overall metabolic frequency profile is less uniform, reflecting a decline in metabolic information uncertainty. **(B)** Conceptualized diagram presenting interpreted patterns of organizations of specialized metabolites according for plant defense theory predictions using information theory descriptors as axes. OD theory predicts that herbivore attack elicits increases in metabolites of defensive value, thereby increasing the RDPI of the metabolome and increasing δj, as particular metabolites with high defensive value are enhanced, thus resulting in high Si. Simultaneously, Hj decreases, as metabolite frequency profiles are reorganized towards a lower metabolic information uncertainty due to the enhancement of herbivory-induced state-specific metabolites. In contrast, MT theory predicts that herbivore attack results in non-directional changes in the metabolome, increasing Hj as an indicator of increased metabolic information uncertainty, no change in δj and creating a random distribution of Si. Using the above information theory descriptors, we additionally propose a mixed model, the Optimal Moving Target (OMT) in which both Hj and δj increase in response to herbivore attack, and metabolites with high defensive value increase in Si, while others exhibit random Si distributions.

Metabolome specialization is measured as the average degree of specificity of individual MS/MS spectra. Consequently, an increase in abundance of certain groups of MS/MSs after herbivore elicitation translate into an increase in profile inducibility, the RDPI index, as well as specialization, the δj index, as more specialized metabolites are produced, generating high Si indexes, whereas a decrease in Hj diversity index reflects that either fewer MS/MSs are being produced or that the profile frequency distribution is changing towards less uniformity, simultaneously reducing its overall uncertainty. With the Si index calculation, it was possible to highlight which MS/MSs are specific to certain herbivory elicitations and reciprocally, which are relatively irresponsive to the elicitation, a key metric that allows MT and OD predictions to be distinguished.

### Predictions of plant defense theories reformulated in the axes of information theory descriptors

Using the information theory descriptors, we interpret the OD theory to predict that the herbivory-induced changes in specialized metabolites from the uninduced constitutive state will result in: (*i*) an increase in metabolome specialization (δj index) driven by increases in the metabolic specificity (Si index) of certain groups of specialized metabolites with high defense value; and (*ii*) a decrease in metabolome diversity (Hj index) caused by changes in the metabolic frequency profiles to more leptokurtic distributions. At the individual metabolite level, an orderly Si distribution in which metabolites increase Si values according to their defense values is expected (Fig. 1B). Along this line, we interpret the MT theory to predict that elicitation will result in: (*i*) a decrease in δj index as a result of non-directional changes in metabolites; and (*ii*) an increase in the Hj index caused by increasing levels of metabolic uncertainty or randomness, the generalized form of diversity which can be quantified by Shannon entropy. As for the metabolic composition, the MT theory would predict a random distribution of Si. Considering that some metabolites are specific to certain conditions but nonspecific to others and their defense value is context-dependent, we additionally propose a mixed defense model in which δj and Hj increase in both directions following the Si distribution that only certain groups of metabolites which possess high defensive values will specifically increase in Si, while others will have a random distribution (Fig. 1B).

### Information theory-analyzed reconfigurations of leaf metabolomes by insect feeding follow OD theory predictions

To test the reformulated predictions of the defense theories in the axes of information theory descriptors, we reared larvae of either the specialist (Ms) or a generalist (Sl) herbivore on leaves of rosette-stage *N. attenuata* plants (Fig. 2A). Using MS/MS analysis, we retrieved 599 non-redundant MS/MS spectra from methanolic extracts of leaf tissues collected after caterpillar feeding (data file S1). Visualizing reconfigurations of the information content in MS/MS profiles using the RDPI, Hj and δj indices revealed intriguing patterns (Fig. 2B). A general trend was a time-dependent, with increases in all degrees of metabolic reorganizations as described by the information descriptors as caterpillars continuously fed on leaves: 72 h after herbivore feeding, RDPI was strongly enhanced; Hj was significantly decreased compared with undamaged controls as a result of an increase specialization of the metabolic profile as quantified by the δj index. This clear trend was in agreement with the predictions of OD theory and inconsistent with the main predictions of MT theory which posits that stochastic (non-directional) changes of metabolite levels are used as a defensive camouflage (Fig. 1B). Surprisingly, direct feeding by the two herbivores, albeit differing in their oral secretion elicitor contents as well as feeding behaviors (*41*), resulted in similar directional changes in Hj and δj at both 24 and 72 h harvests. The only discrepancy occurred in RDPI at 72 h for which Sl feeding elicited a greater overall metabolic inducibility compared to that elicited from Ms feeding.

**Fig. 2.**
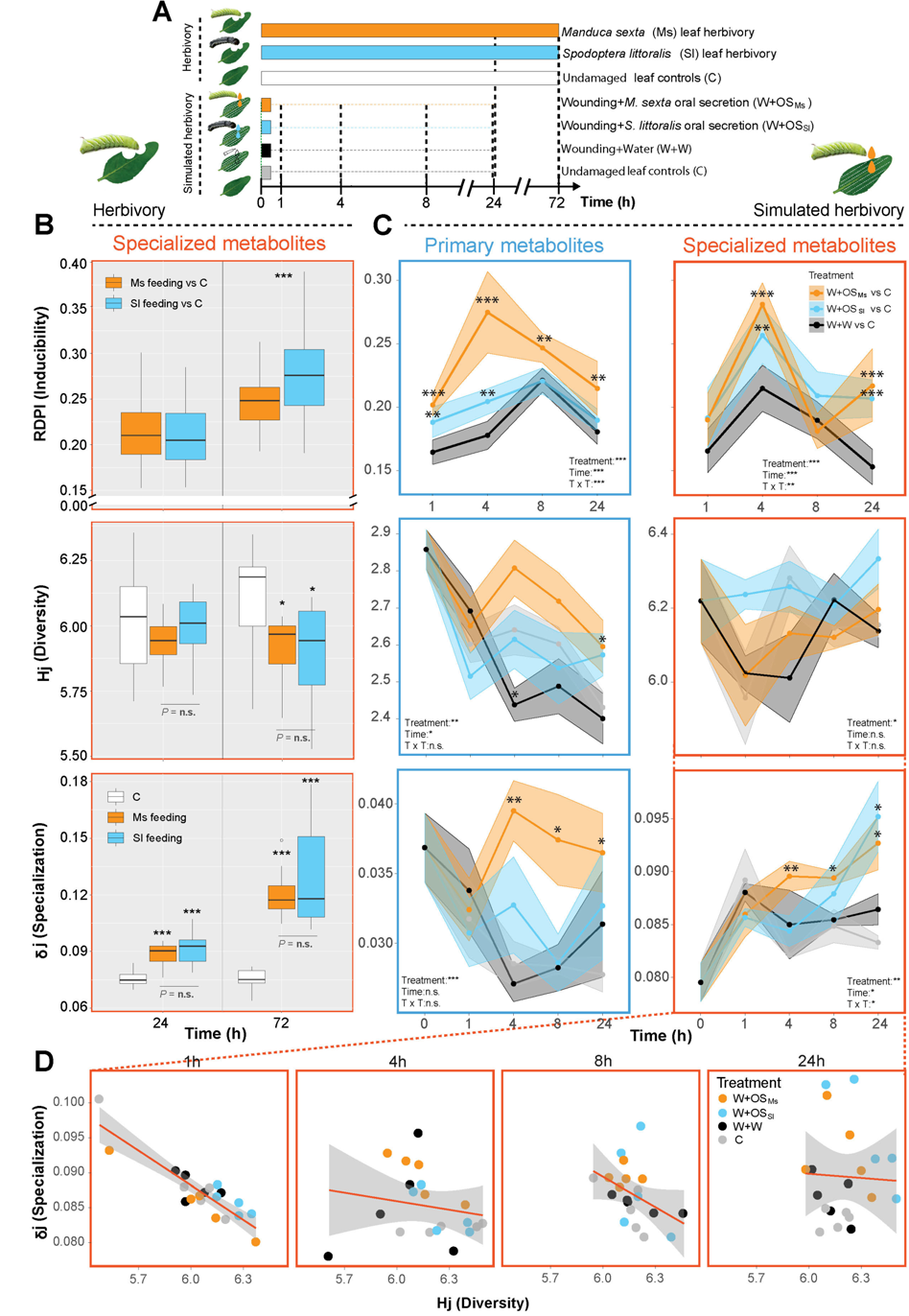
Insect feeding and simulated herbivory reprograms herbivore-specific trajectories of leaf specialized and primary metabolites. (A) Scheme of the experimental design for direct herbivore feeding and simulated herbivory experiments. *Nicotiana attenuata* leaves were fed on by a generalist (*Spodoptera littoralis*: Sl) or specialist (*Manduca sexta*: Ms) herbivore and compared with undamaged control leaves, whereas for simulated herbivory, standardized leaf positions were treating with puncture wounds plus oral secretions (OS) of Ms (W+OS_Ms_) and Sl (W+OS_Sl_) larvae or water (W+W) and compared to undamaged leaves as controls (C). Different color bars indicate different treatment types and black dashed lines indicate sampling times at which treated leaves were harvested. **(B)** Reprogramming of specialized metabolite accumulation in leaves damaged directly by herbivore feeding. Quantification of specialized metabolite plasticity in which inducibility (RDPI), diversity (Hj index) and specialization (δj index) were calculated from the abundance of compound-spectrum-based MS features derived from 599 MS/MSs. RDPI was calculated by comparing data from herbivore-attacked leaves with undamaged controls. Direct herbivore feeding resulted in a temporal increase in RDPI and increased specialization (δj), while diversity (Hj) decreased, which is fully consistent with predictions of the OD but not MT, theory. Asterisks indicate significant differences between treatments (student’s *t*-tests on pairwise differences, \**P* < 0.05, \*\**P* < 0.01, \*\*\**P* < 0.001). n.s., not significant. **(C)** Kinetic analysis of responses to simulated herbivory, which eliminates the confounding inconsistency in tissue loss and timing of the feeding by different insect herbivores, recapitulates the metabolic information remodeling events revealed by direct herbivore feeding. The temporal resolution of the induced responses facilitated by the simulated herbivory treatment reveals herbivore-specific remodeling of the leaf metabolic information landscape. Primary and specialized metabolite inducibility were calculated based on 38 amino acids, organic acids and sugars for primary metabolites (blue box) and 443 MS/MSs for specialized metabolites (red box), respectively. Ribbons refer to 95% confidence intervals. Asterisks indicate significant differences between treatments (Two-way ANOVA followed by *post hoc* multiple comparisons, \**P* < 0.05, \*\**P* < 0.01, \*\*\**P* < 0.001) **(D)** Scatterplot of diversity and specialization of specialized metabolite profiles in which distributions of replicated samples of different treatments analyzed by linear regression models are presented for the indicated time points after the treatment to reveal the changes in metabolic plasticity in high temporal resolution.

To explore if these metabolome-level herbivory-induced reconfigurations are reflected in changes at the level of individual metabolites, we first amortized the vast phytochemical knowledge of *N. attenuata* leaves by focusing on metabolites with proven anti-herbivory functions. Phenolamides are hydroxycinnamic-polyamine conjugates that accumulate during insect herbivory and are known to decrease insect performance (*42, 43*). We retrieved the precursors of corresponding MS/MSs and plotted their accumulation kinetics (fig. S1). As expected, phenolic derivatives such as chlorogenic acid (CGA) and rutin which are not directly involved in anti-herbivory defense were down-regulated after herbivory. In contrast, phenolamides were highly enhanced by herbivory. Interestingly, the continuous feeding by the two herbivores resulted in almost identical elicitation profiles of phenolamides, a pattern which was particularly apparent for the *de novo* synthesized phenolamides. The same phenomenon was observed when exploring the 17-hydroxygeranyllinalool diterpene glycosides (17-HGL-DTGs) pathway that produces abundant acyclic diterpenes with potent anti-herbivore functions (*44*), in which similar expression profiles were triggered by Ms and Sl feeding (fig. S2).

### Simulated herbivory recapitulates directionality and species-specificity of leaf metabolome changes

Possible drawbacks of direct herbivore feeding experiments are differences in leaf consumption rates and timing of feeding making it difficult to disentangle wounding and herbivore-species specific effects induced by herbivory. To better resolve the herbivore species specificity of induced leaf metabolic responses, we mimicked Ms and Sl larval feeding by immediately applying freshly collected oral secretions (OS_Ms_ and OS_Sl_) to standardized puncture wounds (W) in leaves at consistent leaf positions (*45*). This procedure, referred to as W+OS treatment, standardizes the elicitation by precisely timing the initiation of herbivory elicited responses without confounding effects of differences in tissue loss (*46*) (Fig. 2A). Using the MS/MS analytical and computational pipeline, we retrieved 443 MS/MS spectra (data file S1) which largely overlapped with those previously assembled from the direct feeding experiment. In parallel, we quantified levels of 38 amino acids, organic acids and sugars to investigate dynamics of central carbon metabolism. An information theory analysis of this additional MS/MS data-set revealed similar trends in the reprogramming of the leaf specialized metabolome by simulated herbivory as compared to direct herbivore feeding experiment data-set (Fig. 2C). Briefly, metabolic kinetics visualized in a two dimensional space using Hj and δj as coordinates were consistent with a directional increase over time of metabolome specialization in response to simulated herbivory treatments (Fig. 2D). In particular, OS_Ms_ elicited a more significant increase in metabolome specialization at 4 h than did the OS_Sl_ treatment, with main effects detected at the level of primary metabolism intermediates (Fig. 2C). Pathway-level mapping of primary metabolism kinetics further highlighted OS_Ms_-specific increases in amino acids serving as precursors for defensive specialized metabolites, a pattern which contrasts with that of most central organic acid metabolism that exhibited much more conserved responses between OS_Ms_ and OS_Sl_ (fig. S3). To better interpret this pattern, we further monitored metabolic accumulation kinetics of previously discussed phenolamide and 17-HGL-DTG pathways. Interestingly, the herbivore OS-specific induction translated into differential reconfiguration patterns within phenolamide metabolism (fig. S4). Phenolamides containing coumaroyl and caffeoyl moieties were preferentially induced by OS_Sl_ elicitation, while OS_Ms_ triggered a specific induction of feruloyl conjugates. For the 17-HGL-DTG pathway, an OS differential elicitation effect was detected for the downstream malonylated and di-malonylated products (fig. S5).

### An early priming of transcriptome specialization underlies metabolome specialization

The temporal increase of specialization in metabolome elicited by herbivory is likely to translate from coordinated increases in transcriptome specialization. However, a tight correlation of gene and metabolite expression patterns is not always expected, given that certain genes can be constantly transcribed while metabolites accumulate to high levels with long post-stress relaxation times, and reciprocally, although metabolic genes are highly expressed, the corresponding intermediate metabolites are rapidly consumed by subsequent biochemical reactions and may remain at trace levels (*47, 48*). We next investigated OS-elicited transcriptome plasticity using a time-course microarray dataset in which simulated herbivory treatment using OS_Ms_ to leaves of rosette-stage *N. attenuata* plants; the sampling kinetics largely overlapped with those used in the present metabolomics study (*49*). Compared with metabolome reconfigurations in which metabolic plasticity particularly increased over time, we observed a transient burst of transcription in leaves induced by Ms in which transcriptome inducibility (RDPI) and specialization (δj) were strongly enhanced at 1 h, whereas diversity (Hj) was strongly decreased at this time point, followed by a relaxation of transcriptome specialization (fig. S6). Metabolic gene families such as P450, glycosyltransferases and BAHD acyltransferases involved in the assembly of specialized metabolites from building blocks derived from primary metabolism followed the early high specialization pattern described above. The phenylpropanoid pathway was analyzed as a case study, and this analysis confirmed that core genes in phenolamide metabolism were highly OS-induced and tightly co-reconfigured in their expression patterns during herbivory compared to those of unelicited plants. Transcription factor *MYB8* and *PAL1*, *PAL2*, *C4H* and *4CL* structural genes in the upstream part of this pathway exhibited early priming of transcriptions. Acyltransferases such as *AT1*, *DH29* and *CV86* which function as the final assembly of phenolamides exhibited prolonged up-regulated patterns (fig. S6). The above observations suggest that the early priming of transcriptome specialization and the late enhancement of metabolome specialization are coupled patterns, likely as a result of synchronized regulatory systems that launch robust defense responses.

### Phytohormone signaling shapes herbivore-specific changes in the information content of leaf metabolic profiles

Reconfigurations in phytohormonal signaling act as regulatory layers that integrate herbivory information to reprogram a plant’s physiology. We measured accumulation kinetics for key phytohormonal classes after herbivory simulation and visualized temporal coexpression among these (Pearson correlation coefficient (PCC) of >0.4) (Fig. 3A). As expected, phytohormones that are biosynthetically related were linked within the phytohormone coexpression networks. Furthermore, metabolic specificity (Si index) was mapped onto this network to highlight phytohormones that were differentially induced by the different treatments. Two dominant regions of herbivory-specific responses were mapped: one within the jasmonate cluster in which JA, its bioactive form JA-Ile and other JA derivatives exhibited the highest Si scores, and another one was for ethylene (ET). Gibberellins exhibited only moderate increases of herbivore specificity, while other phytohormones such as cytokinins, auxins and abscisic acid exhibited low specificity to the herbivore elicitation. Strong specificity indices for JAs essentially translated from the amplification of peaking values of JA derivatives by OS application (W+OS) compared with W+W alone. Surprisingly, OS_Ms_ and OS_Sl_, which are known to differ in their elicitor contents (*41*), induced similar accumulations of JA and JA-Ile. Ethylene was specifically and strongly induced by OS_Ms_ application, contrasting with OS_Sl_ which did not amplify the basal wound response (Fig. 3B).

**Fig. 3.**
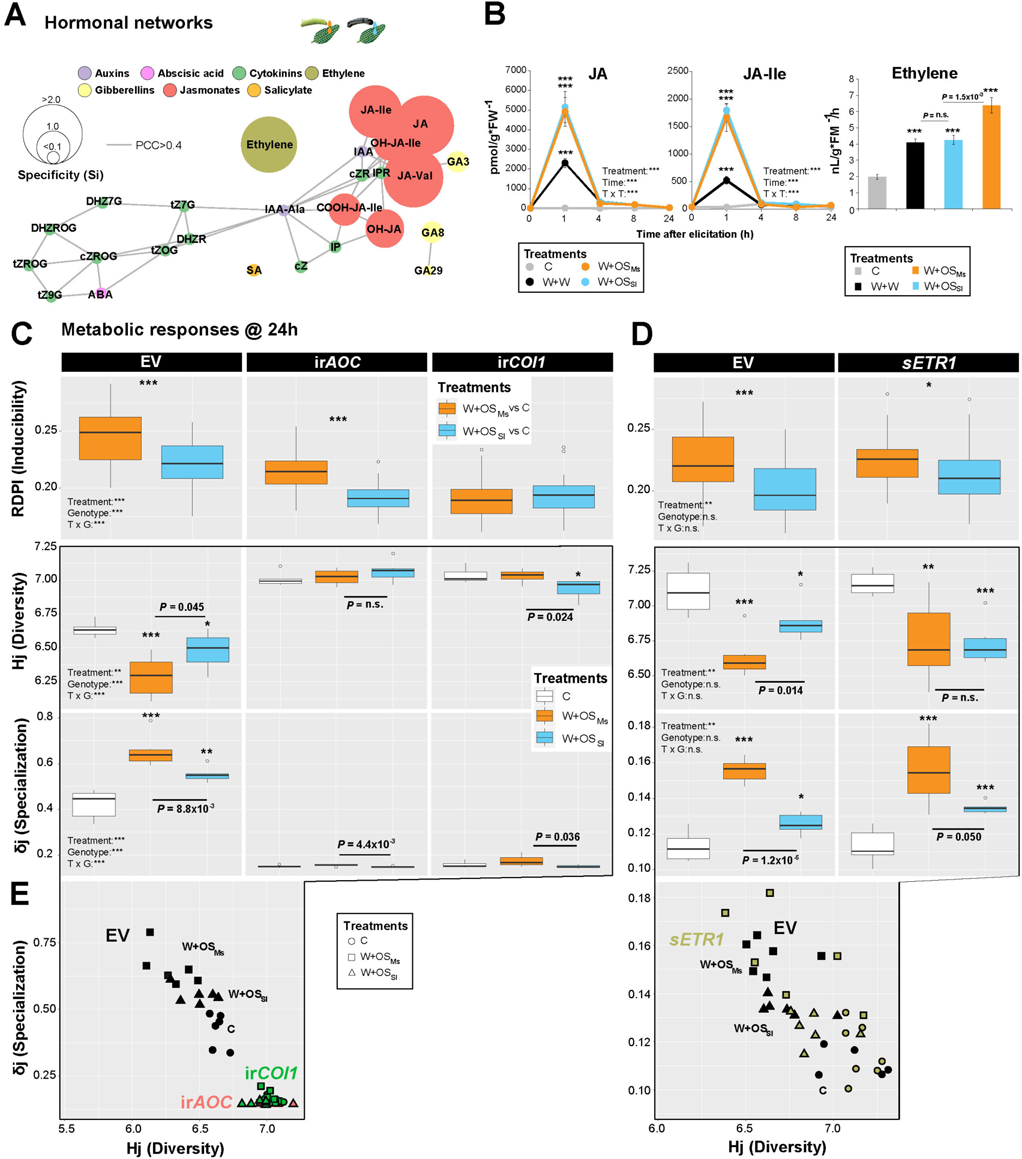
Phytohormone signaling shapes the herbivore-specific metabolic information content of elicited leaves. (**A**) Phytohormone coexpression network analysis which was constructed using Pearson correlation coefficient (PCC) calculations of the accumulation kinetics of key phytohormonal classes after herbivory simulation. Nodes represent individual phytohormone and node size indicates Si indices of phytohormone specificity among treatments. Different colors indicate different phytohormone classes. (**B**) Jasmonate (JA and JA-Ile) and ethylene accumulation patterns in leaves elicited by different treatments indicated with different colors, apricot, W+OS_Ms_; blue, W+OS_Sl_; black, W+W; grey, C (Controls). Information theory analysis of (**C**) 697 MS/MSs (data file S1) in jasmonate biosynthesis and perception-impaired lines (ir*AOC* and ir*COI1*) and of (**D**) 585 MS/MSs in the ethylene signaling-impaired *ETR1* line as well as in empty vector (EV) control plants elicited by the two simulated herbivory treatments, demonstrate that jasmonates function as key determinants of metabolic plasticity whereas ethylene acts as modifier of herbivore-specific responses. Asterisks indicate significant differences between treatments (Two-way ANOVA followed by *post hoc* multiple comparisons, \**P* < 0.05, \*\**P* < 0.01, \*\*\**P* < 0.001). **(E)** Scatterplots of diversity against specialization in jasmonate- and ethylene-impaired transgenic lines. Colors indicate different transgenic lines and linear regression models were calculated for each genotype. Symbols denote for the different treatments, triangle, W+OS_Sl_; rectangular, W+OS_Ms_; circle, C.

Next, we used *N. attenuata* transgenic lines modified in key steps in JA and ET biosynthesis (ir*AOC* and ir*ACO*) and perception (ir*COI1* and *sETR1*) to analyze the relative contribution of these two phytohormones to herbivory-induced metabolic reprogramming. Consistent with the previous experiments, we confirmed a general decrease in Hj index concomitant to an increase in δj index as a result of herbivore-OS elicitation in empty vector (EV) (Fig. 3C-D) and that OS_Ms_-elicited responses were more pronounced than those triggered by OS_Sl_. Specific deregulations were visualized using biplots with Hj and δj as coordinates (Fig. 3E). The most obvious trend was that herbivory-elicited changes in metabolome diversity and specialization were almost fully erased in JA signaling deficient lines (Fig. 3C). In contrast, silenced ET perception in *sETR1* plants, while having a much lower impact on the overall magnitude of herbivory-metabolic changes than JA signaling, attenuated differences in Hj and δj indices between OS_Ms_- and OS_Sl_-elicitations (Fig. 3D and fig S7). This suggested that ET signaling serves as the fine-tuner of the herbivore species-specific metabolic responses in addition to the core function of JA signaling. Consistent with this fine-tuning function, overall metabolome inducibility was not altered in *sETR1* plants. Similar trends were observed using ET deficient ir*ACO* plants (fig. S7).

### Exploring MS/MS herbivore-specific associations using MS/MS structural analysis

To pinpoint specialized metabolites with significant contributions to herbivore species-specific responses and whose production was fine-tuned by ET signaling, we employed a previously developed structural MS/MS approach. This approach relies on a biclustering method to the *de novo* inference of metabolic families from MS/MS fragment (NDP) and neutral loss (NL)-based similarity scoring. The MS/MS dataset constructed from the analysis of ET transgenic lines resulted in 585 MS/MSs (data file S1) which was resolved by biclustering into 7 main MS/MS modules (M) (Fig. 4A). Some of these modules are densely populated with previously characterized specialized metabolites: for instance, M1, 2, 3, 4 and 7 were enriched with various phenolic derivatives, flavonoid glycosides, acyl sugars and 17-HGL-DTGs, respectively. Furthermore, metabolic specificity information (Si index) was also calculated for individual metabolite in each module for which the Si distributions were visualized. Briefly, MS/MS spectra that exhibited high herbivory-elicited and genotypic specificity were characterized by high Si values with kurtosis statistics showing right-tailed leptokurtic distributions. One such leptokurtic distribution was detected for M1 within which phenolamides exhibited the highest Si scores (Fig. 4B). Previously mentioned herbivory-inducible 17-HGL-DTGs within M7 exhibited medium Si scores indicative of a moderate differential regulation by the two OS types. In contrast, mostly constitutively-produced specialized metabolites, such as rutin, chlorogenic acid and acyl sugars were among the lowest Si scores. To better explore structural intricacies among specialized metabolites and their Si distributions, molecular networks were constructed for each of the modules (Fig. 4B). One critical prediction of OD theory (summarized in Fig. 1B) is that the reorganization of specialized metabolites after herbivory should lead to an unidirectional change in metabolites that have high defensive values, particularly through increases in their specificity, a pattern contrasting to the random distribution of defensive metabolites predicted by the MT theory. Most phenolic derivatives clustered in M1 have been functionally associated with decreases in insect performance (*42*). When comparing Si kurtosis distributions within M1 metabolites between induced and constitutive leaves of empty vector (EV) control plants at 24 h, we observed a clear remodeling of kurtosis to heavily right-tailed leptokurtic distributions, indicating that many of the corresponding metabolites in M1 significantly contributed to the metabolome specialization occurring after insect herbivory (Fig. 4C). Specific increases of Si values were exclusively detected for defensive phenolamides but not for other phenolics and yet unknown metabolites coexisting within this module, a specialization pattern which is in agreement with predictions of OD theory. To test whether this specialization of the phenolamide profile translated from OS-specific ET induction, we plotted metabolite Si indices with differential expression values elicited between OS_Ms_ and OS_Sl_ in EV and *sETR1* genotypes (Fig. 4D). In *sETR1*, divergences in phenolamide induction between OS_Ms_ and OS_Sl_ were largely attenuated. The biclustering approach was also applied on MS/MS data collected in JA-deficient lines to infer main MS/MS modules associated to the JA-regulated metabolic specialization (fig. S8).

**Fig. 4.**
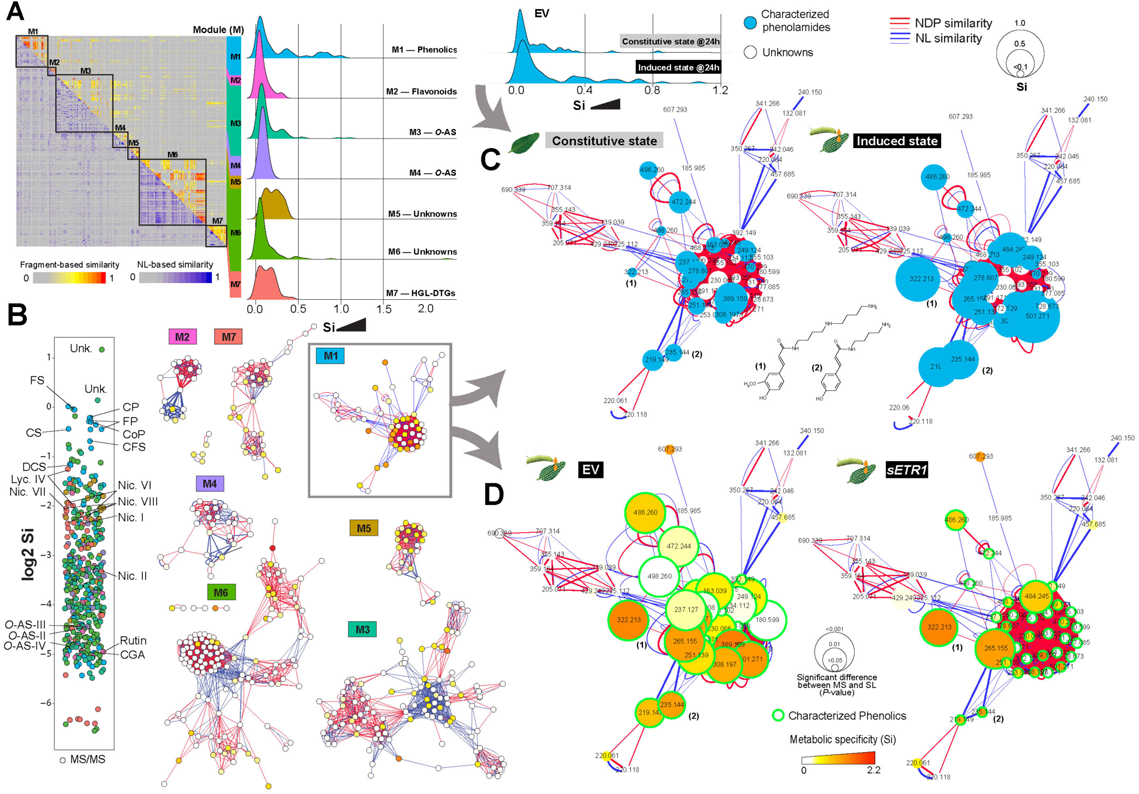
Structural classification of MS/MSs coupled with metabolic specificity information highlights the rearrangement of metabolic specificity distributions after herbivory and reveals a phenolic-enriched module contributing most to the insect species-specific responses. (**A**) Biclustering analysis to classify 585 MS/MSs in ethylene signaling impaired genotypes as well as empty vector (EV) controls according to structural similarities. The analysis used two scoring methods: one based on shared fragments (NDP similarity) among spectra, whereas the other scored shared common neutral losses (NL similarity) among spectra. The biclustering, which favors clustering based on iterative alignments of spectra based on the two scoring methods, produces large modules (M) with structurally related MS/MSs. Some of these modules were congruent with known compound families, whereas others were composed of yet unknown or poorly characterized metabolites. Distributions of metabolic specificity (Si) of the whole dataset are depicted next to each module in which module enrichment is shown. (**B**) Molecular networks constructed for each module. Nodes represent MS/MSs and edges represent similarity values based on the NDP (red) and NL (blue) scoring types. The distribution of metabolic specificity index Si of each MS/MS was ranked and visualized for all modules (left panel) and further mapped onto the molecular networks for each module (right panel). **(C)** A comparative view of Si distribution rearrangement in M1 between constitutive state (undamaged leaf controls) and induced state (after simulated herbivory) in EV at 24h. The corresponding molecular networks (NDP and NL similarity cutoff > 0.6) of M1 are presented in which phenolic compounds that are known to function as defense against herbivory are highlighted in blue and the Si values are shown as node sizes. The analysis illustrates that metabolites with defensive properties were particularly enhanced in their metabolic specificity Si after herbivory elicitation, a pattern consistent with the predictions of OD theory, while inconsistent with the predictions of MT theory. (**D**) Molecular networks of M1 comparing EV with the ethylene perception impaired line, *ETR1*. The green circled nodes represent annotated phenolic compounds according to the structural classification pipeline and the node sizes indicate the degree of significant differences (*P*-value) between W+OS_Ms_ and W+OS_Sl_ treatments. The herbivore-specific differentiation of phenolic compound accumulation is abolished when ethylene perception is impaired. CP, *N*-caffeoyl-putrescine; CS, *N*-caffeoyl-spermidine; FP, *N*-feruloyl-putrescine; FS, *N*-feruloyl-spermidine; CoP, *N’,N’’*-coumaroyl-putrescine; DCS, *N’, N’’*-dicaffeoyl-spermidine; CFS, *N’,N’’*-caffeoyl, feruloyl-spermidine; CGA, Chlorogenic acid; Lyc., lyciumoside; Nic., nicotianoside; *O*-AS, *O*-acyl sugars; Unk., unknown.

### Natural variation in jasmonate bursts underlie intra-specific variations in herbivory-induced metabolome specialization

We further extended the analysis from a single *N. attenuata* genotype to natural populations in which intense intra-specific variations in herbivory-induced JA levels and specialized metabolites levels have been previously described (*36*). Using this dataset which covers 43 accessions consisting of 123 individual *N. attenuata* plants derived from seeds collected at different native habitat in Utah, Nevada, Arizona and California (Fig. 5A), we calculated their metabolome diversity and specialization shaped by OS_Ms_ elicitation (Fig. 5B). Consistent with the previous study, we observed a wide range of metabolic variations along Hj and δj axes, indicating that accessions differ dramatically in the plasticity of their metabolic responses to herbivory (Fig. 5C). The significant positive correlation (PCC=0.62, *p*-value=1.7×10^-14^) between population-level metabolic diversity (here referred to as population-level β-diversity) and specialization is depicted in the 2D scatter space in Fig. 5C. This positive correlation implies that herbivory-elicited remodeling of metabolomes in populations results from synchronous modulations of metabolome β-diversity and specialization. Such organization is reminiscent of previous observation made on the dynamic range of variation in herbivory-induced levels of jasmonates and that extreme values are maintained within single populations (*36, 50*). By testing population-level correlations between Hj and δj with JA and JA-Ile, we found significant positive correlations between JAs and both metabolome β-diversity and specialization indices (Fig. 5C). This suggested that the herbivory-induced heterogeneity in the elicitation of JAs detected at the population level is likely a key metabolic polymorphism in response to insect herbivory.

**Fig. 5.**
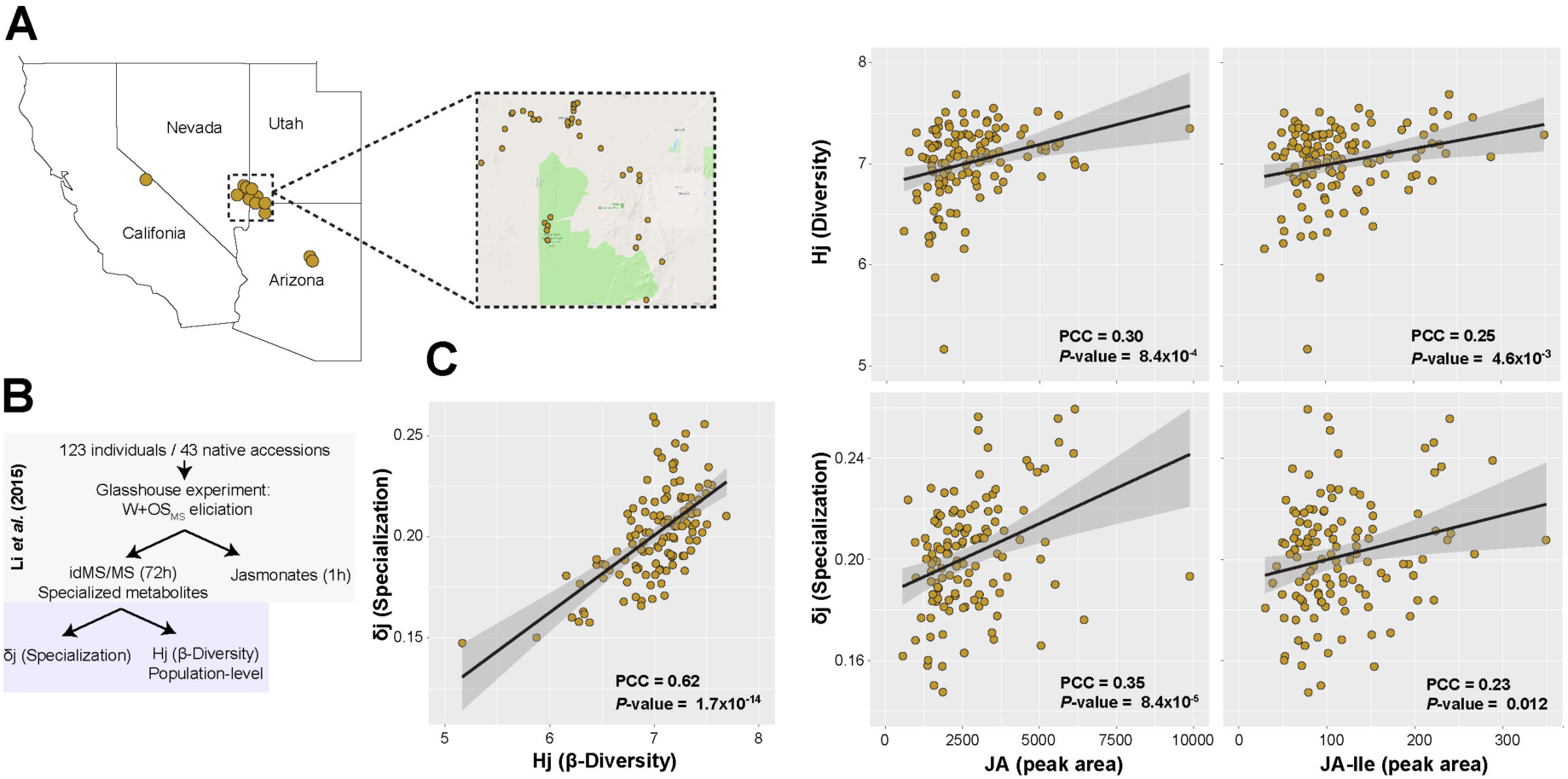
Population-level OS-elicited metabolic diversity and specialization are positively correlated with jasmonate signaling. (A) The geographic location of the *N. attenuata* seed collection sites of 43 accessions consisting of 123 individuals in Utah, Nevada, Arizona, and California. A close-up for the collection sites in Utah is presented**. (B)** Workflow of the extraction of population-level metabolic diversity and specialization. **(C)** Scatter plot of diversity (Hj) versus specialization (δj) extracted from metabolic profiles of leaves of 123 plants harvested at 72h after W+OS_Ms_ treatment. At the intraspecific level, β-diversity is positively correlated with specialization. At the population level, both metabolic diversity and specialization are positively correlated with jasmonates (JA and JA-Ile). Density plots are shown adjacent to the scatter plots. Pcc, pearson correlation coefficient.

### Exploring assignments of OS-elicited responses in closely-related *Nicotiana* species reveals species-specific coordination of specialized metabolites with jasmonate plasticity

Previous research has shown that *Nicotiana* species largely differ in the type and relative reliance on inducible vs constitutive metabolic defenses. These variations in anti-herbivore signaling and defenses are thought to be modulated by insect population pressures, plants’ life cycles, as well as the costs of defense production within the ecological niche in which a given plant species grows. We examined the consistency of herbivory-induced reconfigurations of leaf metabolomes in 6 *Nicotiana* species, native to North and South America and closely-related to *N. attenuata*, namely, *Nicotiana pauciflora, N. miersii, N. attenuata, N. acuminata, N. linearis, N. spegazzinii and N. obtusifolia* (*51*) (Fig. 6A). 6 of these species, including the well-characterized species *N. attenuata*, are annuals from the *Petunioides* clade, while *N. obtusifolia* is a perennial plant from the sister clade *Trigonophyllae* (*52*). These 7 species were subsequently subjected to W+W, W+OS_Ms_ and W+OS_Sl_ induction to study species-level metabolic reconfigurations to insect herbivory. Using the biclustering approach, we identified 9 modules of 939 MS/MSs (data file S1). The MS/MS compositions that were reconfigured by different treatments differed greatly in different modules among species (fig. S9). Visualizing Hj (here referred to as species-level γ-diversity) and δj revealed that species clustered as very distinct groups in the metabolic space, species-level demarcation effect being frequently more prominent than the elicitation effects.

**Fig. 6.**
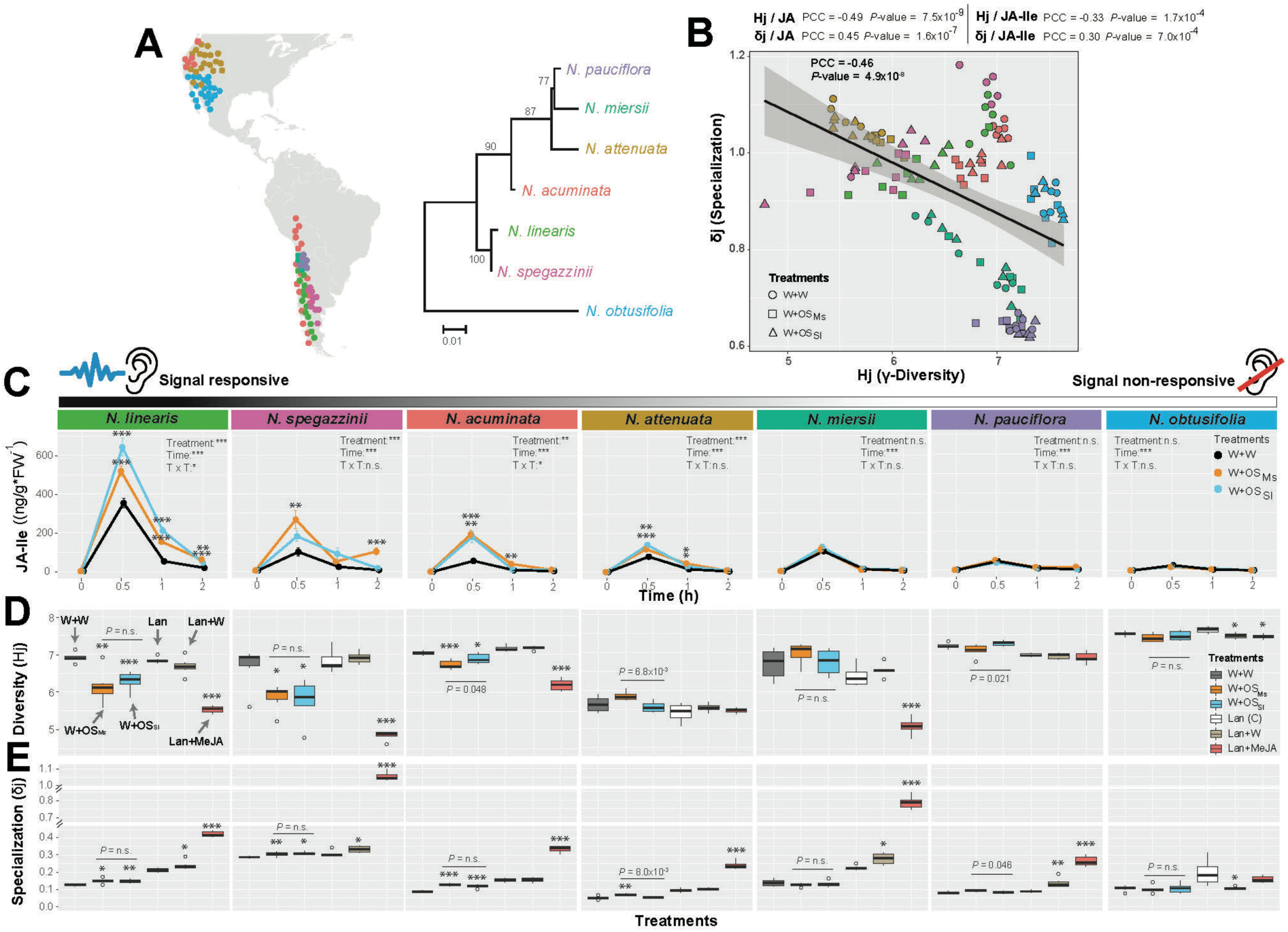
OS-elicited specialized metabolite responses in closely related *Nicotiana* species reveal species-specific trajectories that are highly correlated with plasticity in jasmonate signaling. (**A**) The phylogenetic tree and geographic distribution of seven closely related *Nicotiana* species. The phylogenetic tree was constructed using nuclear glutamine synthetase sequences obtained from Clarkson *et al*. 2010 (52). The evolutionary history was inferred by using the Maximum Likelihood method based on the Tamura-Nei model. Bootstrap values supporting the tree are shown next to the branches. The geographic distributions of the species were from Goodspeed (51). Different colors indicate different species. (**B**) Information theory analysis of 939 MS/MSs constructed using the seven closely-related *Nicotiana* species sampled at 72h after treatments. At the species level, diversity, which is defined as γ-diversity, was negatively correlated with specialization. A linear regression model was drawn across species. Species-level correlational analysis of metabolic diversity and specialization with jasmonate (JA and JA-Ile) accumulation is shown above the scatter plot (PCC, pearson correlation coefficient). γ-diversity was negatively correlated with jasmonates whereas specialization was positively correlated with jasmonates. Detailed correlational analysis is shown in fig. S10. Colors indicate different species and symbols indicate different treatments, triangle, W+OS_Sl_; rectangular, W+OS_Ms_; circle, C. (**C**) JA-Ile kinetics of the seven species that are ranked according to their maximum JA-Ile bursts and plasticity in response to OS elicitation. Species such as *N. linearis* and *N. spegazzinii* are highly plastic in their jasmonate signaling in response to simulated herbivory, while others such as *N. obtusifolia* are insensitive to simulated herbivory. Asterisks indicate significant differences between treatments (Two-way ANOVA followed by *post hoc* multiple comparisons, \**P* < 0.05, \*\**P* < 0.01, \*\*\**P* < 0.001). Scatter plots of diversity to specialization for each species in **(D)** simulated herbivory and **(E)** methyl jasmonate (MeJA) applications with linear regression models, demonstrating that species-specific plasticity of metabolic reconfigurations in response to herbivory are highly coordinated with species-specific plasticity in jasmonate signaling.

Exceptions were *N. linearis* and *N. spegazzinii* which exhibited broad dynamic ranges of elicitation effects (Fig. 6B). In contrast, species such as *N. pauciflora* and *N. obtusifolia* exhibited less pronounced metabolic responses to treatments but greater metabolome diversities. The species-specific distributions of induced metabolic responses resulted in a significant negative correlation between specialization and γ-diversity (PCC = −0.46, *p*-value=4.9×10^-8^). Variations in OS-induced levels of JAs were positively correlated with metabolome specialization, but negatively correlated with metabolic γ-diversity exhibited by each species (Fig. 6B and fig. S10). Notably, species colloquially referred to as “signal responsive” ones in Fig. 6C, such as *N. linearis*, *N. spegazzinii*, *N. acuminata* and *N. attenuata* elicited remarkable OS-specific JA-Ile burst at 30 min, while others referred to as “signal non-responsive” ones such as *N. miersii*, *N. pauciflora* and *N. obtusifolia* only showed marginal inductions of JA-Ile and without any OS-specificity (Fig. 6C). At the metabolic level and as previously described for *N. attenuata*, “signal responsive” species exhibited an OS-specific and significant increase δj concomitant with a decrease Hj. This OS-specific elicitation effect was not detected in species classified as “signal non-responsive” ones (Fig. 6, D and E). Remarkably, the OS-specifically elicited metabolites are shared more frequently among “signal responsive” species, which clustered apart from the “signal non-responsive” ones that exhibited less species interdependencies (fig. S11). This result suggested that the OS-specific induction of JAs and the OS-specific reconfigurations of downstream metabolomes were coupled patterns at the species level. We next investigated whether these coupling patterns were constrained by JA availability using exogenous JA application by treating plants with lanolin paste containing MeJA, which is rapidly de-esterified to JA in the plant’s cytoplasm. Interestingly, we found the same trend of graded changes from signal-responsive to signal-non-responsive species induced by the continuous supply of JA (Fig. 6, D and E). Briefly, the MeJA treatment strongly reprogramed the metabolomes of *N. linearis*, *N. spegazzinii*, *N. acuminate*, *N. attenuata* and *N. miersii* towards pronounced increases of δj and decreases of Hj. *N. pauciflora* only showed an increase in δj but not Hj. Remarkably, *N. obtusifolia* which was previously shown barely capable of inducing JAs, barely responded to MeJA treatment. These results suggested that JA production or signal transduction is physiologically constrained in the “signal non-responsive” species. To test this hypothesis, we investigated the transcriptomes of 4 of the species (*N. attenuata*, *N. miersii*, *N. pauciflora* and *N. obtusifolia*) induced by W+W, W+OS_Ms_ and W+OS_Sl_ (*53*). Consistent with the pattern of metabolome reconfigurations, species were well-separated in the transcriptomic space with *N. attenuata* exhibited the highest OS-induced RDPI while *N. obtusifolia* being the lowest (Fig. 7A). However, the transcriptome diversity induced by *N. obtusifolia* was found to be the lowest among the 4 species, a pattern opposite to the highest metabolome diversity of *N. obtusifolia* previously shown among 7 species. Previous research has revealed that a group of genes involved in early defense signaling, including JA signaling, accounted for the specificity of herbivore associated elicitors induced early defense responses within *Nicotiana* species (*53*). Comparing the JA signaling pathway among the 4 species revealed interesting patterns (Fig. 7B). A majority of genes in this pathway, such as *AOC*, *OPR3*, *ACX* and *COI1*, exhibited comparably high induction levels among the 4 species. However, a key gene, *JAR4*, which converts JA to its bioactive form JA-Ile accumulated transcripts at very low levels specifically in *N. miersii*, *N. pauciflora* and *N. obtusifolia*. Additionally, transcripts of another gene, *AOS,* were not detected only in *N. obtusifolia*. These changes in gene expression are likely to be responsible for the low JA production in the “signal non-responsive” species and the extreme phenotype of constrained induction for *N. obtusifolia*.

**Fig. 7.**
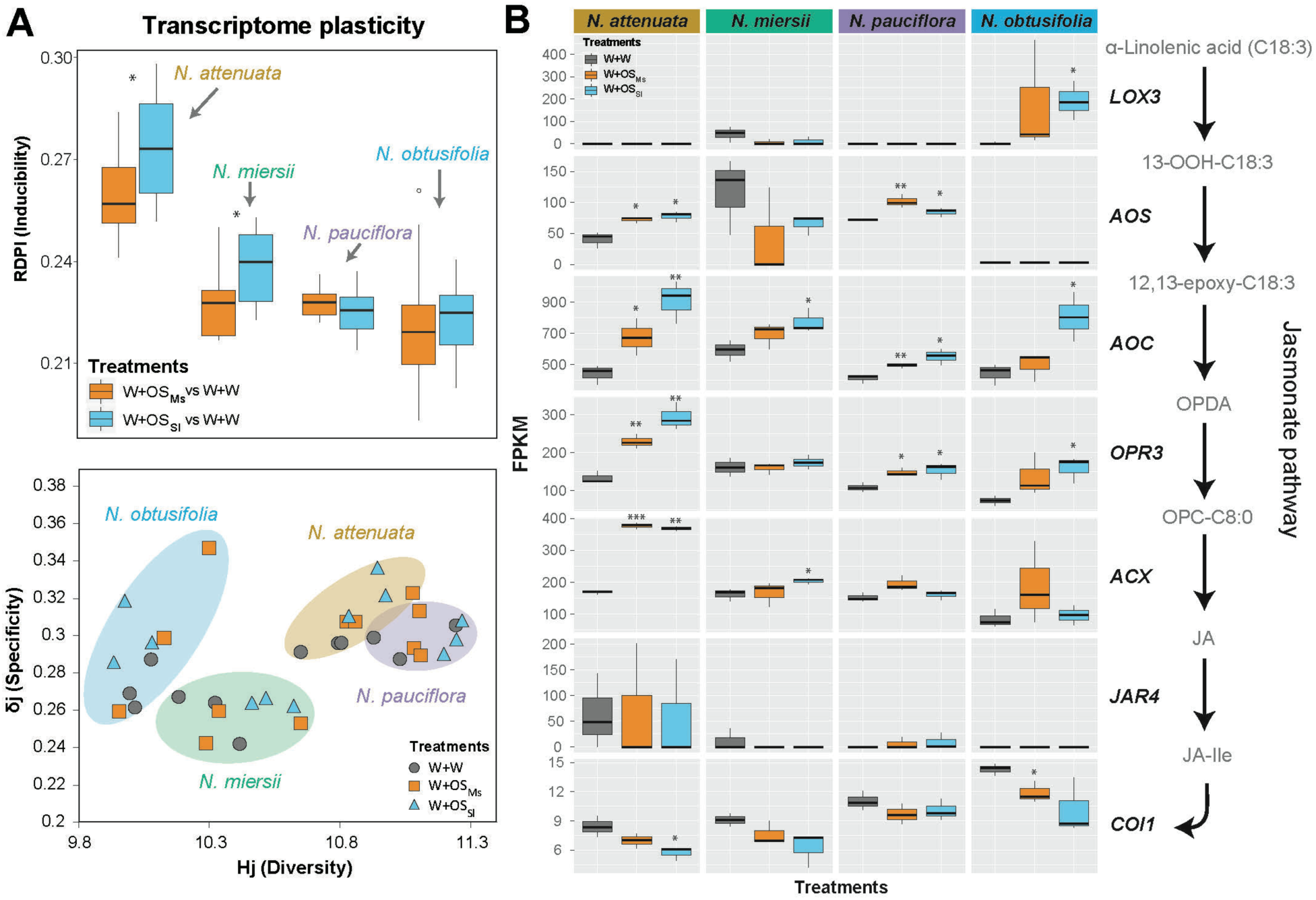
OS-elicited transcriptome responses in closely related *Nicotiana* species highlight key genes in the jasmonate signaling pathway responsible for the plant species-specific responses to herbivory. (**A**) Information theory analysis of the reprogramming of early transcriptomic responses of 4 closely related *Nicotiana* species sampled 30 min after herbivory induction. RDPI was calculated by comparing herbivore OS-elicited leaves with wounding controls. Colors indicate different species and symbols indicate different treatments. (**B**) Gene expression analysis in the JA signaling pathway among the 4 species. A simplified JA pathway is shown next to box plots. Different colors indicate different treatments. Asterisks indicate significant differences compared to wounding controls (\**P* < 0.05, \*\**P* < 0.01, \*\*\**P* < 0.001).

### Impact of species-specific induced responses on herbivore resistance

Finally in this last section, we examine how the insect species-specific remodeling of metabolomes of different plant species contributed to herbivore resistance. Previous research has highlighted that *Nicotiana spp.* differed greatly in their induced resistance to Ms and Sl larval performance (*54*). Here we investigated how this pattern was linked to their metabolic plasticity. Using the above mentioned 4 *Nicotiana* species and testing the correlation between herbivory-induced metabolome diversity and specialization and the plants’ resistance to Ms and Sl, we found that the resistance to the generalist Sl was positively correlated with both diversity and specialization, whereas the resistance to the specialist Ms showed weaker correlations with specialization and was not significant correlated with diversity (fig. S12). Intriguingly for the resistance to Sl, both *N. attenuata* and *N. obtusifolia*, previously shown to differ greatly in their responses to herbivore induction at both the level of JA signaling and metabolome plasticity, showed similarly high levels of resistance to Sl.

## Discussion

Over the past six decades, plant defense theories have provided theoretical frameworks from which researchers have made predictions about the evolution and function of the considerable diversity of the specialized metabolites of plants. Most of these theories have not followed the normal procedures of strong inference (*55*) presenting critical predictions posed at the same levels of analysis (*3*) which would allow the field to move forward when the tests of critical predictions allow a particular theory to be supported, while rejecting others (*56*). Instead, new theories made predictions at different levels of analysis, adding new descriptive layers of consideration (*56*). However, two theories, MT and OD, posed at the functional level of analysis, could be readily interpreted as making critical predictions regarding herbivore-induced changes in specialized metabolism: OD theory predicted that the changes in specialized metabolism “space” would be highly directional, favoring metabolites with high defensive value, while the MT theory posited that the changes would be non-directional and randomly positioned in the metabolic space. Here, we used innovations in computational MS-based metabolomics and deconvoluted MS analyses in the common currencies of information theory descriptors to test the critical predictions that distinguish the two theories. Information theory has been applied in many fields, particularly in ecology in the context of biodiversity and trophic flow studies (*7*). But, as far as we know, this is the first application to describe plants’ metabolic information space and address ecological questions regarding the function of temporal metabolic changes in response to environmental cues. In particular, the power of the present approach lies in its ability to compare patterns within and among plant species to examine how herbivory has shaped plant metabolism at different scales in the evolutionary hierarchy, from micro-evolutionary patterns within species to among-species macro-evolutionary patterns.

Decomposing the information content of leaf specialized metabolite profiles in terms of specificity, diversity and inducibility, revealed that herbivore feeding activated idiosyncratic metabolic rearrangements. Surprisingly, we observed that the resulting metabolic landscapes, as described by the implemented information theory indices, largely overlapped after attack by the two herbivores, Sl, a nocturnal feeding generalist and Ms, a Solanaceae specialist, despite their distinct feeding behaviors and content in fatty acid-amino acid conjugates (FAC) elicitors in their OS (*41*). Simulated herbivory treatments by treating standardized puncture wounds with herbivore OS revealed similar trends. This standardized procedure of mimicking a plant’s responses to herbivore attack removes confounding factors caused by variations in herbivore feeding behaviors that result in different amounts of damage occurring at different times (*45*).

FACs which are known to be the major elicitors in OS_Ms_ that induce JA and other phytohormonal responses in *N. attenuata* are hundreds of times lower in OS_Sl_ (*41*). However, OS_Sl_ elicited similar levels of JA accumulations compared to OS_Ms_. The JA response in *N. attenuata* was previously shown to be very sensitive to OS_Ms_ in which FACs can retain its activity even when diluted 1:1000 with water (*57*). It is thus likely that the FACs in OS_Sl_, albeit low, were sufficient to elicit a full induction of JA burst compared to OS_Ms_. Previous studies have shown that porin-like proteins (*58*) and oligosaccharides (*59*) can serve as molecular cues in OS_Sl_ that trigger plant defense responses. However, it is still unclear whether these elicitors in OS_Sl_ were responsible for the JA accumulations observed in the current study.

While few studies have described differential metabolic fingerprints induced by different herbivores or exogenous JA or SA applications (*60, 61*), none have conducted a systemic exploration of herbivore species-specific perturbations in a plant’s phytohormonal network and its holistic consequences for specialized metabolic profiles. The present analysis further confirmed that intrinsic hormonal network connections with other phytohormones beyond JAs shape the specificity of herbivory-induced metabolic reconfigurations. In particular, we detected a dramatically larger elicitation of ET by OS_Ms_ than by OS_Sl_. This pattern is consistent with the greater FACs content in OS_Ms_ which is necessary and sufficient to elicit ET burst (*62*). The herbivore-specific induction of ET was shown to ultimately modulate the metabolome response, most notably for the herbivore-specific *de novo* accumulation of phenolamides as revealed by transgenic manipulations of ET biosynthetic (*ACO*) and perception (*ETR1*) genes. This synchronized ET modulation of specialized metabolome was further revealed when herbivores continuously fed on leaves (fig. S13). ET has previously been shown to fine-tune the JA-induced nicotine accumulation by regulating putrescine *N*-methyltransferases (*63*). Besides ET’s signaling functions, investments into polyamine-based phenolamides could possibly be modulated via metabolic flux shunt, due to the competition for *S*-adenosyl-l-methionine involved as common intermediate in both ET and polyamine biosynthetic pathway.

Strong focus towards specific metabolic classes due to the sheer number of specialized metabolites of unknown structures have long precluded rigorous assessments of temporal shifts in metabolic diversity following biotic interactions (*64*). A central result from the present information theory analysis based unbiased metabolite-derived MS/MS spectra acquisition was that herbivore feeding or simulated herbivory consistently lowers the overall metabolic diversity of leaf metabolomes while increasing their degree of specialization. Remarkably, features contributing most (with high Si values) to this greater metabolome specialization were specialized metabolites with previously characterized anti-herbivory functions. This pattern is consistent with the predictions of the OD theory, but not with MT predictions regarding the stochasticity of the metabolome reprogramming. However, the data was also consistent with predictions of a mixed model (OMT, Fig. 1B), as other uncharacterized metabolites possessing yet unknown defensive functions may still follow a random Si distribution. One striking pattern further captured by this study was that different levels of evolutionary organizations, from micro-evolutionary levels (single plants and *Nicotiana* populations) to larger evolutionary scales (closely-related *Nicotiana* species), differed dramatically in their capacities to “optimally defend” against herbivores. Moore et al. (*28*) and Kessler (*1*) have independently proposed to transpose the three functional levels of biodiversity originally distinguished by Whittaker (*65*) to constitutive and induced temporal changes in chemodiversity, these authors did not outline procedures either for large-scale metabolome data acquisition or for how metabolic diversity could be calculated from these data. In the present study, minor adjustments to Whittaker’s functional categorization would consider α-metabolic diversity as the diversity of MS/MS spectra in a given plant, β-metabolic diversity as the fundamental intra-specific metabolic space for a set of populations, while γ-metabolic diversity would be the extension of the analysis to congeneric species.

We observed that herbivory-induced metabolome specialization in *N. attenuata* and variations of metabolome specialization within *N. attenuata* populations and among closely-related *Nicotiana* species were systematically positively correlated with JA signaling. Furthermore, the herbivory-induced metabolic specialization on a single genotype basis was largely abolished when JA signaling was impaired (Fig. 3, C and E). As changes in the metabolic profiles of natural *N. attenuata* populations were mostly quantitative, the amplitude of metabolic β-diversity variations in this analysis was more constrained than that of the specialization index but remained positively correlated with JA signaling. More revealing was the analysis of closely-related *Nicotiana* species as those differed greatly in their biochemical machineries, resulting in qualitatively specialized metabolite profiles. The information theory processing of the captured metabolic profiles revealed a trade-off, exacerbated by the herbivory induction, between metabolic γ-diversity and specialization. JA signaling plays a central role in this trade-off as increases in metabolome specialization which is consistent with the main OD prediction was positively correlated with JA signaling, while JA signaling was negatively correlated with metabolic γ-diversity. These patterns suggested that plant’s optimal defense capacities are largely defined by the JA plasticity at both micro-evolutionary and larger evolutionary scales. Exogenous JA application experiments, to by-pass JA biosynthetic deficiencies, further revealed that closely-related *Nicotiana* species could be distinguished as “signal responsive” and “signal non-responsive” species just as they were by their patterns of JA and metabolome plasticity in response to herbivory induction. “Signal non-responsive” species were physiologically constrained by their inability to produce and respond to endogenous JA, likely due to mutations in a few key genes in the JA signaling pathway (*AOS* and *JAR4* in *N. obtusifolia*) of which transcripts were absent. This result highlights that these interspecific macro-evolutionary patterns may largely be driven by variations in internal hormone perception and responsiveness.

Beyond plant-herbivore interactions, exploring metabolic diversity is relevant to all important theoretical advances in the study of organisms’ adaptations to their environments and complex phenotypic trait evolution. With the increase of data volumes acquired by modern MS instruments, hypothesis testing regarding metabolic diversity can now transcend differences in individual/classes of metabolites and proceed to global analyses revealing unsuspected patterns. In this larger-scale analysis process, an important metaphor is the idea of constructing a meaningful map from which data can be explored. As such, an important outcome of the present combination of unbiased MS/MS metabolomics and information theory is that it provides a simple metric with which to construct a map that allows for the browsing of metabolic diversity at the different taxonomical scales, an essential requirement for the study of micro/macro-evolution and community ecology. At the macro-evolutionary level, core to Ehrlich & Raven’s plant-insect coevolution theory (*66*) is the prediction that interspecific variations in metabolic diversity are responsible for lineage diversification in plants. However in the five decades since this seminal work was published, few tests of this hypothesis have been conducted (*67*), in large part due to the phylogenetic rarity of comparable metabolic characters across distant plant lineages that can be used to anchor targeted analytical measurements. The present information theory-processed MS/MS workflows enable such a taxonomic scale comparison of these macro-evolutionary patterns in specialized metabolism by quantifying MS/MS structural similarities of unknown metabolites, without *a priori* metabolite selection, and translate these MS/MSs into a set of simple statistical indices. The process is analogous to a phylogenetic analysis which quantifies the rate of diversification or character evolution using sequence alignment without *a priori* predictions. At the biochemical level, the screening hypothesis by Jones and Firn (*68*) implies that metabolic diversity is maintained at different hierarchical scales in order to provide the raw material to exapt bioactivities of previously extraneous or alternatively adapted metabolites. Information theory approach provides a framework in which to quantify these evolutionary transitions in metabolite specificity that should occur during metabolite exaptation as part of the proposed evolutionary screening processes: from low to high specificity for exapted metabolites whose bioactivity becomes adaptive for a given environmental context.

In conclusion, during the early days of molecular biology when the seminal plant defense theories were developed, deductive hypothesis-driven methods were widely considered to be the only means of making scientific advances, in large part due to the technical constraints of measuring entire metabolomes. While hypothesis-driven approaches are particularly useful in choosing among alternative causal mechanisms, their ability to advance our understanding of biochemical networks is more limited compared to the computational methods currently available in contemporary data-intensive science. Consequently, theories which made predictions far beyond the reach of the available data could not be fully falsified thereby abrogating the hypothesis formulation/testing cycle through which a research field makes progress (*4*). We foresee that the metabolomics computational workflow presented here could reinvigorate interest in both proximate (how) and ultimate (why) questions regarding metabolic diversity and contribute to a new era of theory-guided data science that revisits the important theories that inspired previous generations.

## Materials and Methods

### Plant Treatment and Sample Preparation

Direct herbivore feeding was conducted by rearing one second instar *M. sexta* or *S. littoralis* larvae on one *N. attenuata* leaf of individual rosette-stage plants, each with 10-plant replicates. Insect larvae were clip-caged and remaining leaf tissues were collected and flash-frozen at 24 and 72 h after infestation, and metabolites were extracted.

Simulated herbivory treatment was conducted in a highly synchronized fashion by producing, with a fabric pattern wheel, three rows of punctures onto each side of the midvein of three fully expanded leaves from plants in the rosette-stage of growth and immediately applying 1:5 diluted *M. sexta* or *S. littoralis* oral secretions (OSs) to the puncture wounds with a gloved finger. One treated leaf was harvested and processed as described above. Primary metabolites and phytohormones were extracted using previously described methods (*69*).

For exogenous JA applications, three petioles of leaves from each of six rosette-stage plants per species were treated with either 20 µl lanolin paste containing 150 µg methyl jasmonate (Lan + MeJA) or with 20 µl lanolin plus wounding treatment (Lan+W) or with 20 µl pure lanolin as control. Leaves were harvested at 72 h after treatment, flash frozen in liquid nitrogen, and stored at −80°C until use.

### Ethylene measurement

Ethylene measurements were conducted noninvasively with a photoacoustic laser spectrometer (INVIVO) as previously described in von Dahl et al. (*70*). For direct herbivore feeding, leaves were excised and transferred to 50 mL bottles where one second instar *M. sexta* or *S. littoralis* larvae was allowed to feed freely on the excised leaf and the headspace ethylene that had accumulated every 2 h was quantified before flushing with ethylene-free air.

### Microarray and RNAseq data analysis

Microarray data were originally published in (*49*) and deposited in the National Center for Biotechnology Information Gene Expression Omnibus database (accession no. GSE30287). Data corresponding to leaves elicited by W+OS_Ms_ treatments as well as undamaged controls, were extracted for the present study. Raw intensities were log2 and baseline transformed and normalized to their 75^th^ percentile using the R software package, prior to statistical analysis. The raw RNAseq data of *Nicotiana* species were retrieved from the NCBI short reads archive (SRA) under the project number PRJNA301787 which was reported by Zhou et al. (*53*) and processed as described in (*71*). Raw data corresponding to W+W, W+ OS_Ms_ and W+OS_Sl_ treatments of *Nicotiana* species were selected for the present study analysis and processed as follows: raw RNA-seq reads were first converted to fastQ format. HISAT2 converted fastQ to sam, and SAMtools converted sam files to sorted bam files. StringTie was used to calculate gene expression as fragments per kilobase of transcript per million reads sequenced (FPKM).

### MS/MS collection, deconvolution and similarity scoring

Data-independent or indiscriminant MS/MS fragmentation analysis (hereafter referred to as MS/MS) was conducted in order to gain structural information on the overall detectable metabolic profile. Injection and UHPLC binary gradient-based separation conditions used for the MS and MS/MS mode analyses were previously described in ref. 11, MS/MS assembly was achieved via correlational analysis between MS1 and MS/MS mass signals for low and high collision energies and newly implemented rules. The correlation analysis for precursor-to-product assignment for was implemented using an R script and rules were implemented using a C# script (https://github.com/MPI-DL/indiscriminant-MS-MS-assembly-pipeline). MS/MS spectra were aligned in a pairwise manner and their similarity calculated according to a standard normalized dot product (NDP) for fragment similarity and a Euclidean distance for shared neutral losses.

### Defining inducibility, diversity, specialization and metabolic specificity using information theory

Metabolic profile inducibility was calculated using Relative Distance Plasticity Index (RDPI) (*40*). Metabolic profile diversity, the Hj index, was calculated using Shannon entropy of MS/MS frequency distribution derived from the abundance of MS/MS precursors. Metabolic profile specialization, the δj index, was measured by the average MS/MS specificity of each of the MS/MS component for a given sample. Metabolic specificity, the Si index, was defined as the expression identity of a given MS/MS regarding frequencies among considered samples.

## Acknowledgments

We thank Wenwu Zhou for helping with the samples, Rishav Ray for the support in assembling the RNAseq dataset and Han Guo and Zachariah Jaramillo for the help in establishing ethylene measurement.’

## Author contributions

D.L. conceived the study, performed the experiments, analyzed the data and wrote the paper; R.H. helped conceive the study; E.G. and I.T.B. conceived the study and wrote the paper.

## Data and materials availability

Microarray data are deposited in National Center for Biotechnology Information Gene Expression Omnibus database under accession no. GSE30287. All raw RNAseq data of *Nicotiana* species are accessible under NCBI short reads archive (SRA) under the project number PRJNA301787. All other data needed to evaluate the conclusions in the paper are available within the main text or supplementary materials.

## Supporting Information

## Materials and Methods

### UHPLC-ESI/qTOF-MS conditions for profile mode analysis and MS/MS data acquisition

An Acclaim column (150×2.1 mm, particle size 2.2 µm) with a 4 mm×4 mm guard column of the same material was used for the analysis. The following binary gradient was used with a Dionex Ultimate 3000 UHPLC system: 0 to 0.5 min, isocratic 90% A (de-ionized water, 0.1% [v/v] acetonitrile and 0.05% formic acid), 10% B (acetonitrile and 0.05% formic acid); 0.5 to 23.5 min, gradient phase to 10% A, 90% B; 23.5 to 25 min, isocratic 10% A, 90% B. Flow rate was 400 µL/min. For all MS analyses, the column eluent was infused into an Impact II (Bruker Daltonics, Bremen, Germany) equipped with quadrupole and time-of-flight analyzers and fitted with an electrospray source operated in positive ionization mode (capillary voltage 4500 V, capillary exit 130 V, dry temperature 200 °C, dry gas flow of 10 L/min).

The concept of the indiscriminant MS/MS approach relies on the fact that the quadrupole is operated with a very large mass isolation window (so that quasi all *m/z* signals are considered for fragmentation). For this, several independent analyses are performed with increasing CID collision energy (CE) values since the Impact II instrument cannot create CE ramping. Briefly, samples were first analyzed by UHPLC-ESI/qTOF-MS using the single MS mode (low fragmentation condition derived from in-source fragmentation) by scanning from *m/z* 50 to 1500 at a repetition rate of 5 Hz. MS/MS analyses were conducted using nitrogen as collision gas and involved independent measurements at the following 4 different collision-induced dissociation voltages: 20, 30, 40 and 50ev. The quadrupole was operated throughout the measurement with the largest mass isolation window, from *m/z* 50 to 1500. This mass range was automatically activated by the operating software of the instrument when the precursor *m/z* and the isolation width are experimentally set to 200 and 0 Da, respectively. Mass fragments were scanned as in the single MS mode. Mass calibration was performed using sodium formate (50 mL isopropanol, 200 µL formic acid, 1 mL 1M NaOH in water). Data files were calibrated post-run on the average spectrum from a given time segment, using the Bruker HPC (high-precision calibration) algorithm. Raw data files were converted to netCDF format using the export function of the Data Analysis v4.0 software (Bruker Daltonics, Bremen, Germany).

### Additional rules for the assembly of compound-specific MS/MS

To reduce false positive errors resulting from spurious correlations from background noise due to the fact that some *m/z* features are only detected in a few samples, we compared data processing results obtained with and without the “fill peaks” function of XCMS (use for background noise correction) and calculated a background noise value from the average correction estimate used by this function to replace “NA” not detected peak intensities. When the “fill peaks” function is used, there still were many “0” intensity values in the dataset which affect the calculation of correlations, and these were replaced with the calculated background value. We also only considered features with intensities that were more than 3 times the background value and considered these as “true peaks”. Only *m/z* signals with at least eight “true peaks” for the samples precursors (MS1) and fragments datasets were considered for PCC calculation. A precursor mass feature is further defined if its intensity across sample significantly correlate with the decreased intensity of the same mass feature subjected to low or high collision energies and that this feature is not annotated as an isotope peak by CAMERA. The correlation analysis was then conducted by calculating all possible precursor-to-product pairs within 3s – estimated retention time window for peak deviation. *m/z* values were only considered as fragments if they were lower than that of the precursor and MS/MS fragmentation occurred in the same sample position within the dataset as the precursor from which it is derived. Based on these two simple rules, we excluded assigned fragments at *m/z* values larger than that of the identified precursor as well as based the sample position for occurrence precursor and assigned fragments. Many in-source-fragmentation-generated mass features produced in the MS1 mode can also be selected as candidate precursors resulting in redundant compound MS/MS. To reduce such data redundancy, we merged spectra if their NDP similarity exceeded 0.6 and they belong to the chromatographic “pcgroup” annotated by CAMERA. Finally, we merged all the 4 collision energy results for precursor-to-fragment associations into a final deconvoluted composite spectrum by choosing the highest intensity peak among all candidate peaks of the same *m/z* value at the different collision energies. This latter processing step is based on the composite spectrum concept and accounts for the different collision energy conditions required to maximize fragmentation possibilities since certain fragments are detected only at specific collision energies.

### Information theory framework for defining metabolome diversity and specialization and metabolic specificity

Metabolome diversity was calculated using Shannon entropy of MS/MS frequency distribution by the following equation as described in Martinez et al., (2008) (*10*):

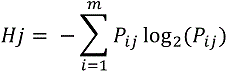

where P_ij_ correspond to relative frequency of the ith MS/MS (i = 1, 2, …, m) in the jth sample (j = 1, 2, …, t).

The average frequency of the i^th^ MS/MS among samples was calculated as:

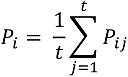

MS/MS specificity was calculated as:

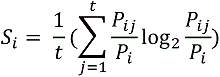

The metabolome specialization δj index was measured for each j^th^ sample, the average of the MS/MS specificities using the following formula:

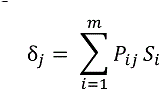

### MS/MS similarity scoring

MS/MS spectra were aligned in a pairwise manner and their similarity calculated according to two scores. First, a standard normalized dot product (NDP), also referred to as cosine correlation method, was used to score fragment similarity among spectra using the following equation:

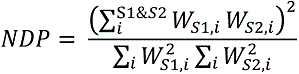

where S1 and S2 correspond, respectively, to spectrum 1 and spectrum 2 and *W*_S1,i_ and *W*_S2,i_ indicate peak intensity-based weights given to *i*^th^common peaks differing by less than 0.01Da between the two spectra. Weights were calculated as follows:

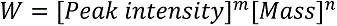

with m = 0.5 and n = 2 as suggested by MassBank (*72*).

A second scoring method involving the analysis of shared neutral losses among individual MS/MS was implemented. For this, we used a list of 52 neutral losses (NLs) commonly encountered during tandem MS fragmentation as well as more specific ones that had been previously annotated for MS/MS spectra of *N. attenuata* secondary metabolite classes (table S1) (*11, 36*). A binary vector of 1 and 0 was created for each MS/MS corresponding to present and absent of certain NL. NL similarity score was calculated for each pair of binary NL vectors based on Euclidean distance similarity.

### MS/MS molecular networking by bi-clustering

To perform this clustering, we used the R package DiffCoEx which is based an extension of the Weighted Gene Coexpression Analysis (WGCNA). Using NDP and NL-scoring matrices for MS/MS spectra, we computed a comparative correlation matrix using DiffCoEx with the parameters of “cutreeDynamic” set to method=“hybrid”, cutHeight = 0.9999, deepSplit = T, minClusterSize = 10. The R source code of DiffCoEx is downloaded from additional file 1 in Tesson et al. (2010) (*73*), the required R WGCNA package can be found at http://www.genetics.ucla.edu/labs/horvath/CoexpressionNetwork/Rpackages/WGCNA.

### Microarray experimental design

Wild type (WT) *N. attenuata* plants were grown in 16 h light/8 h dark cycle for 5 weeks. Rosette leaves of *N. attenuata* plants were wounded with a fabric pattern wheel and the resulting puncture wounds were immediately treated with 20 µL of diluted oral secretions (OS, 1∶5 with distilled water) from the larvae of the specialist herbivore, *M. sexta*. Treated leaves were harvested first at 1 h and then every 4 h after W+OS elicitation for 1 day. Detailed experimental setup is described in (*49*).

### Files

**Data file S1.** XCMS-processed data-sets and deconvoluted MS/MS data-sets

---

**Figure S1.**
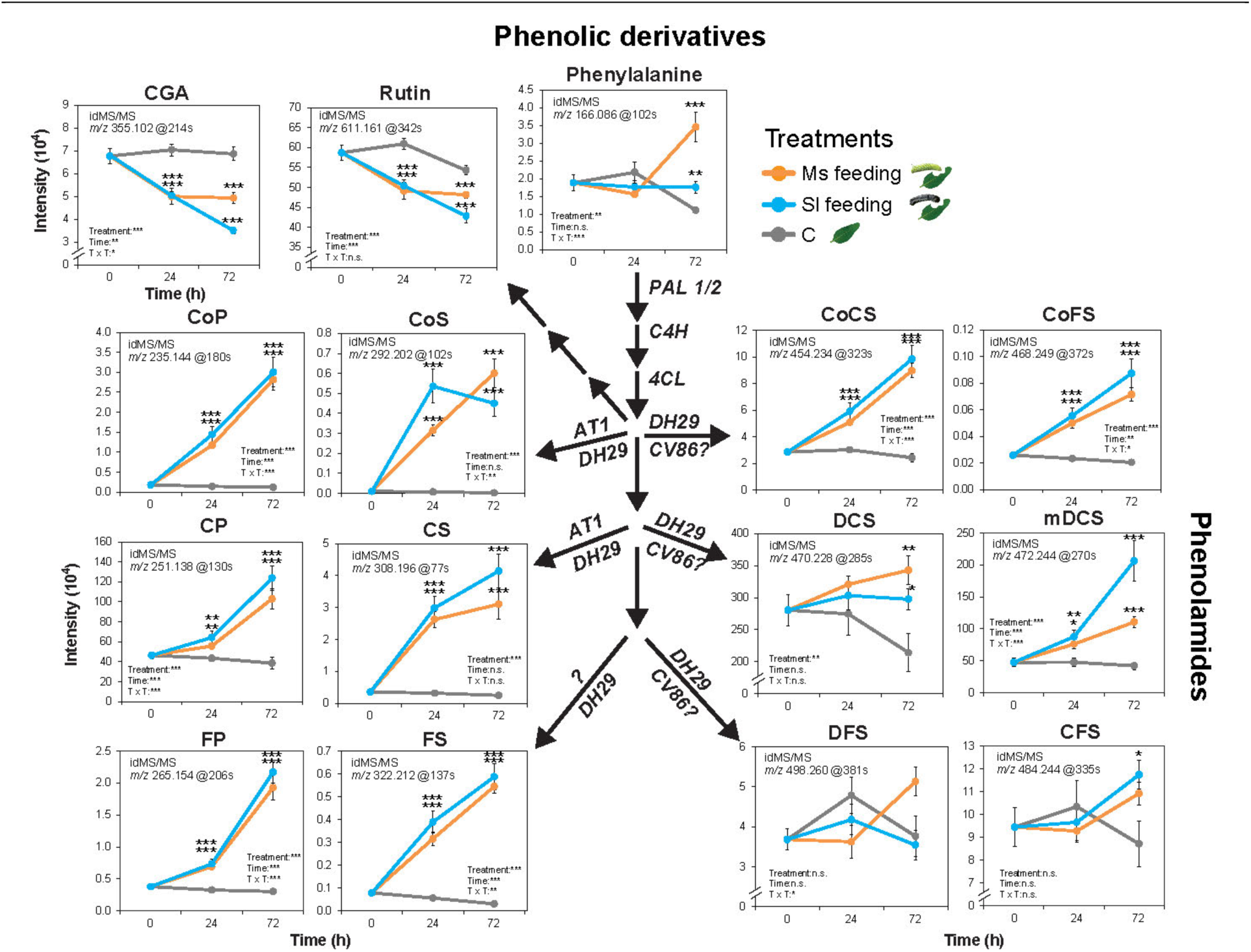
Accumulation of phenolic derivatives, including phenolamides (phenylpropanoid-polyamine conjugates), in *N. attenuata* leaves in response to continuous herbivore feeding. A simplified pathway connects MS/MSs corresponding to major phenylpropanoid-quinate and -polyamine conjugates and illustrates branch-specific differential regulation upon insect feeding. CGA, Chlorogenic acid; CoP, *N*-coumaroyl-putrescine; CoS, *N*-coumaroyl-spermidine; CP, *N*-caffeoyl-putrescine; CS, *N*-caffeoyl-spermidine; FP, *N*-feruloyl-putrescine; FS, *N*-feruloyl-spermidine; CoCS, *N’,N’’*-coumaroyl, caffeoyl-spermidine; CoFS, *N’,N’’*-coumaroyl, feruloyl-spermidine; DCS, *N’, N’’*-dicaffeoyl-spermidine; mDCS, Monohydrated *N’, N’’*-dicaffeoyl-spermidine; DFS, *N’,N’’*-diferuloyl-spermidine. CFS, *N’,N’’*-caffeoyl,feruloyl-spermidine. Asterisks indicate significant differences between treatments (Two-way ANOVA followed by *post hoc* multiple comparisons, *P < 0.05, **P < 0.01, ***P < 0.001). n.s., not significant. Ms, *Manduca sexta*; Sl, *Spodoptera littoralis*; C, control.

**Figure S2.**
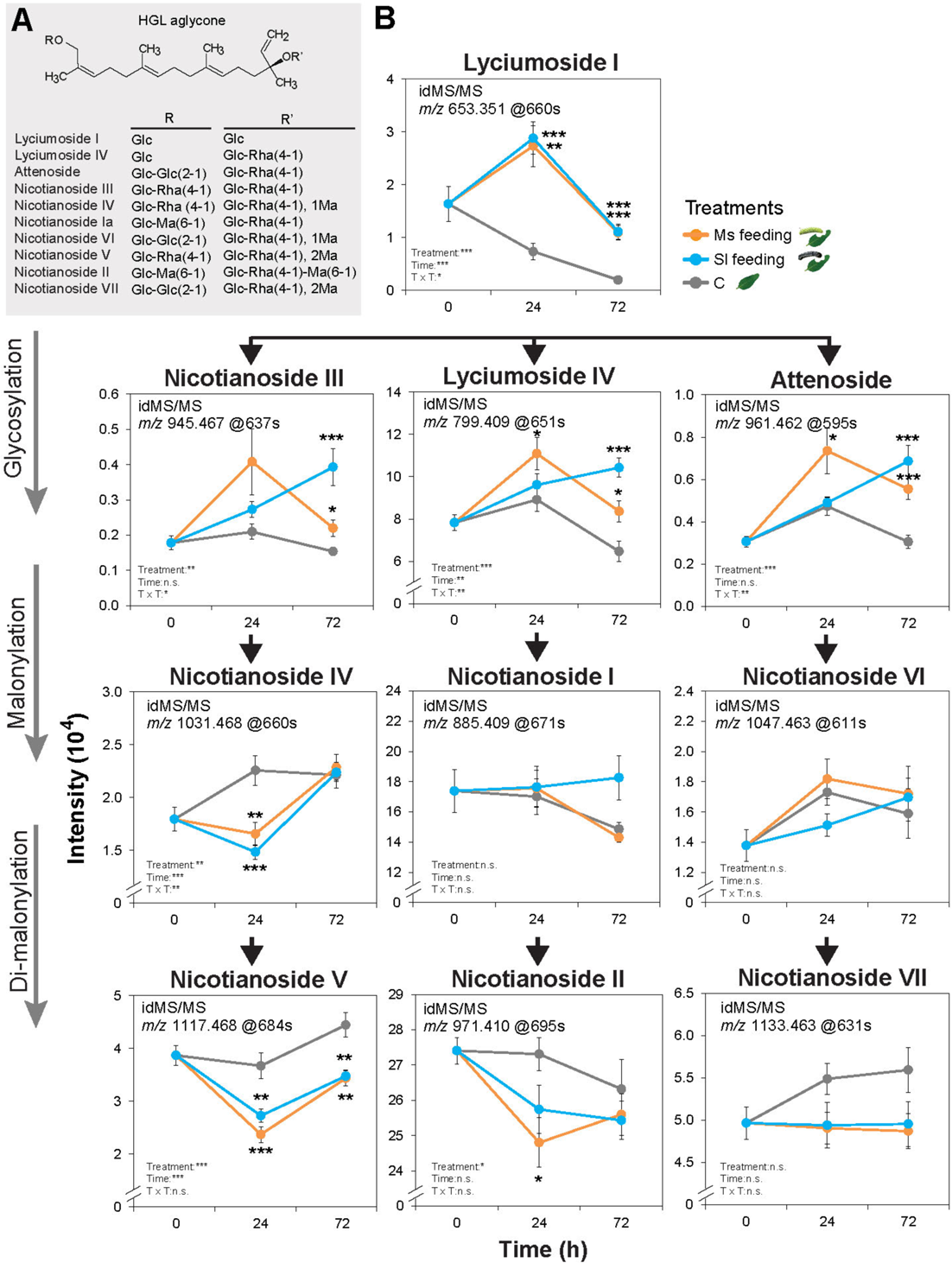
Accumulation of 17-HGL-DTGs in *N. attenuata* leaves in response to continuous herbivore feeding. (**A**) 17-hydroxygeranyllinalool (HGL) diterpene glycosides are abundant secondary metabolites in *N. attenuata* which differ in the number and types of sugar (Glc, glucose; Rha, Rhamnose; with or without malonyl, Ma, groups) decorations added to the acyclic HGL backbone which is characteristic of this compound family. (**B**) Pathway-level illustration of differential patterns of induction of 17-HGL-DTGs by herbivore feeding. MS/MSs corresponding to most abundant HGL-DTGs are presented. Asterisks indicate significant differences between treatments (Two-way ANOVA followed by *post hoc* multiple comparisons, *P < 0.05, **P < 0.01, ***P < 0.001).

**Figure S3.**
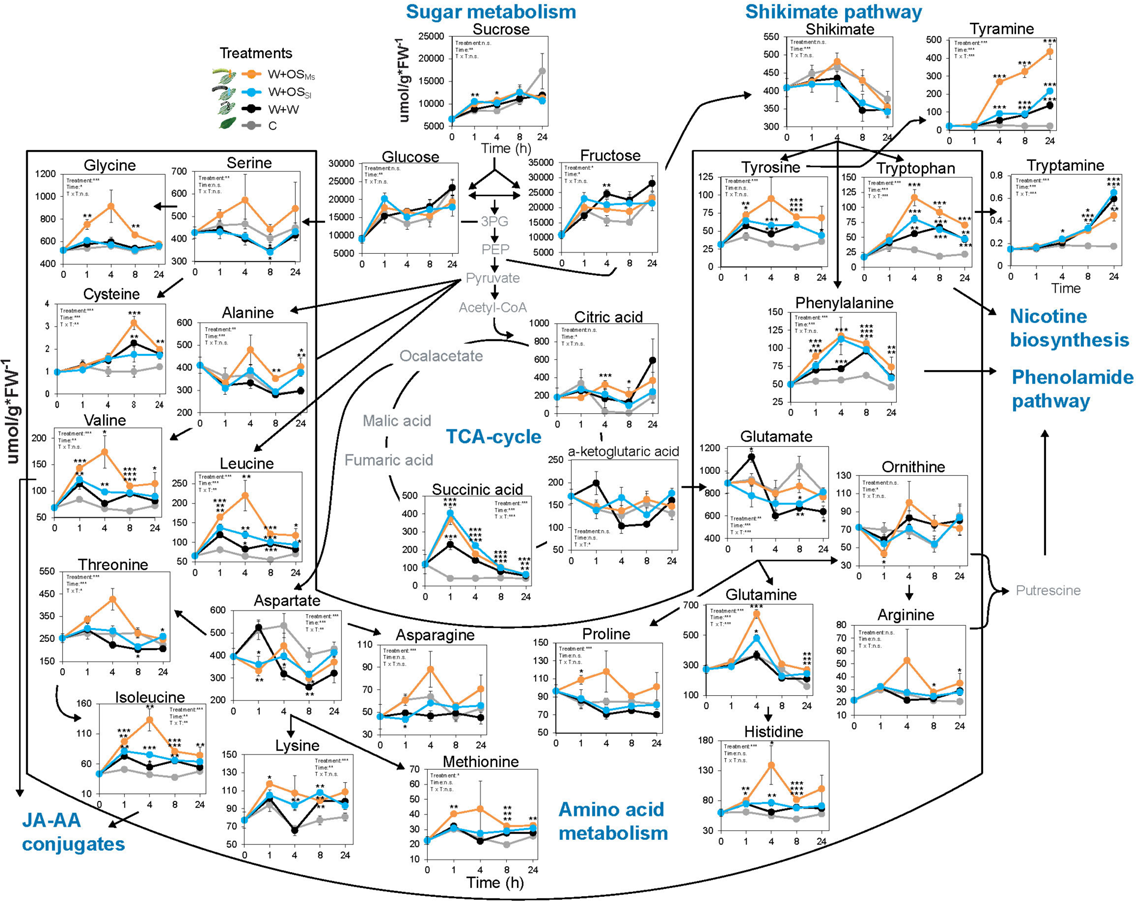
Herbivory-induced reconfigurations of metabolites located within central metabolism. Primary metabolites were measured and quantified using a previously described targeted method (*69*). Compounds that were not included in the analysis are only designated by name and depicted in grey font color. Asterisks indicate significant differences between treatments (Two-way ANOVA followed by *post hoc* multiple comparisons, *P < 0.05, **P < 0.01, ***P < 0.001). For simulated herbivory treatments, standardized leaf positions were wounded with a fabric pattern wheel and the resulting puncture wounds were treated with oral secretions (OS) of Ms (W+OS_Ms_) and Sl (W+OS_Sl_) larvae or water (W+W) and compared to undamaged leaves as controls (C).

**Figure S4.**
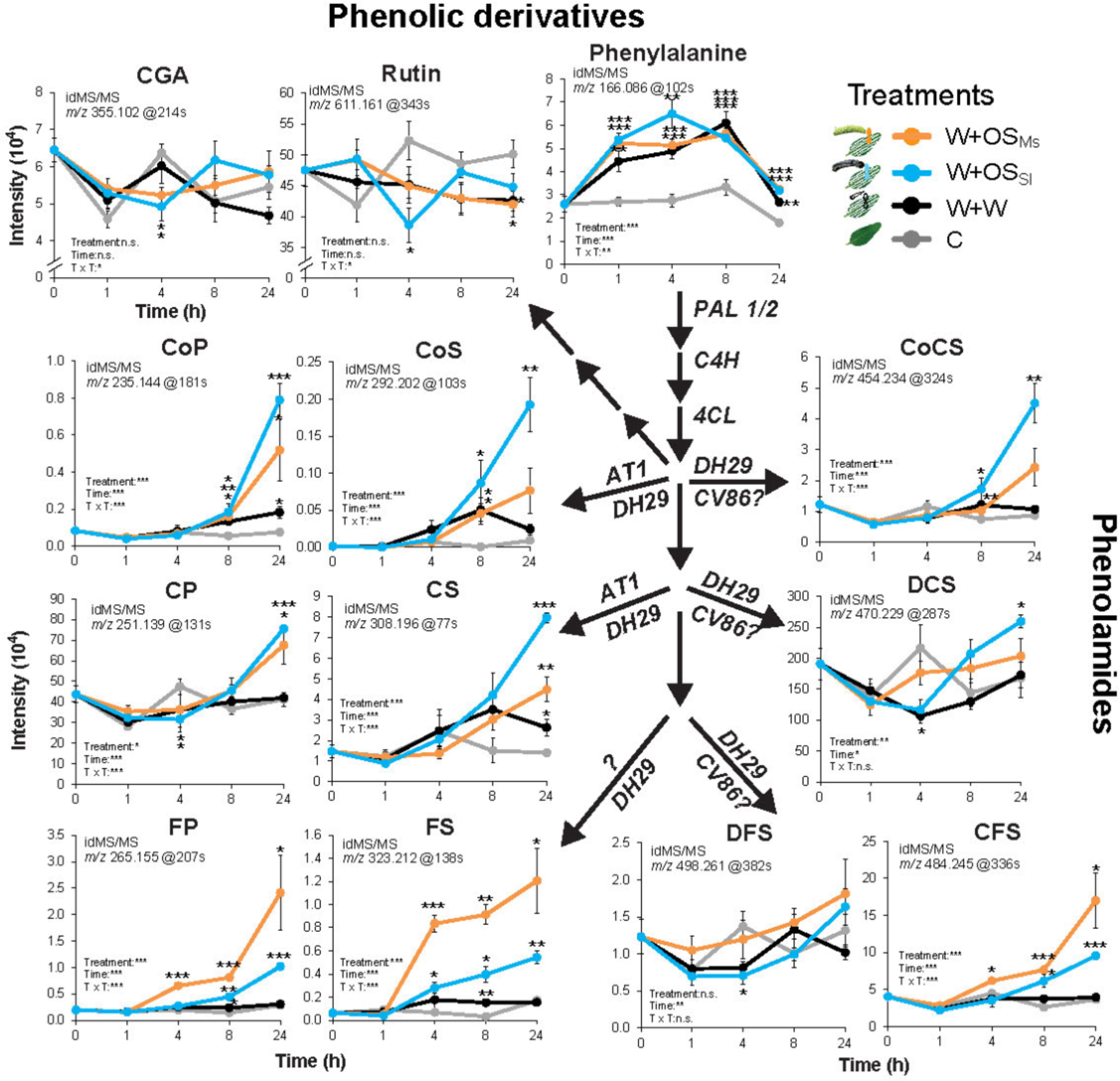
Accumulation of phenolic derivatives, including phenolamides (phenylpropanoid-polyamine conjugates), in *N. attenuata* leaves in response to simulated herbivory. A simplified phenolic pathway with MS/MSs corresponding to major phenylpropanoid-quinate and -polyamine conjugates illustrating metabolic differential accumulation patterns at different pathway branches. Asterisks indicate significant differences between treatments (Two-way ANOVA followed by *post hoc* multiple comparisons, *P < 0.05, **P < 0.01, ***P < 0.001).

**Figure S5.**
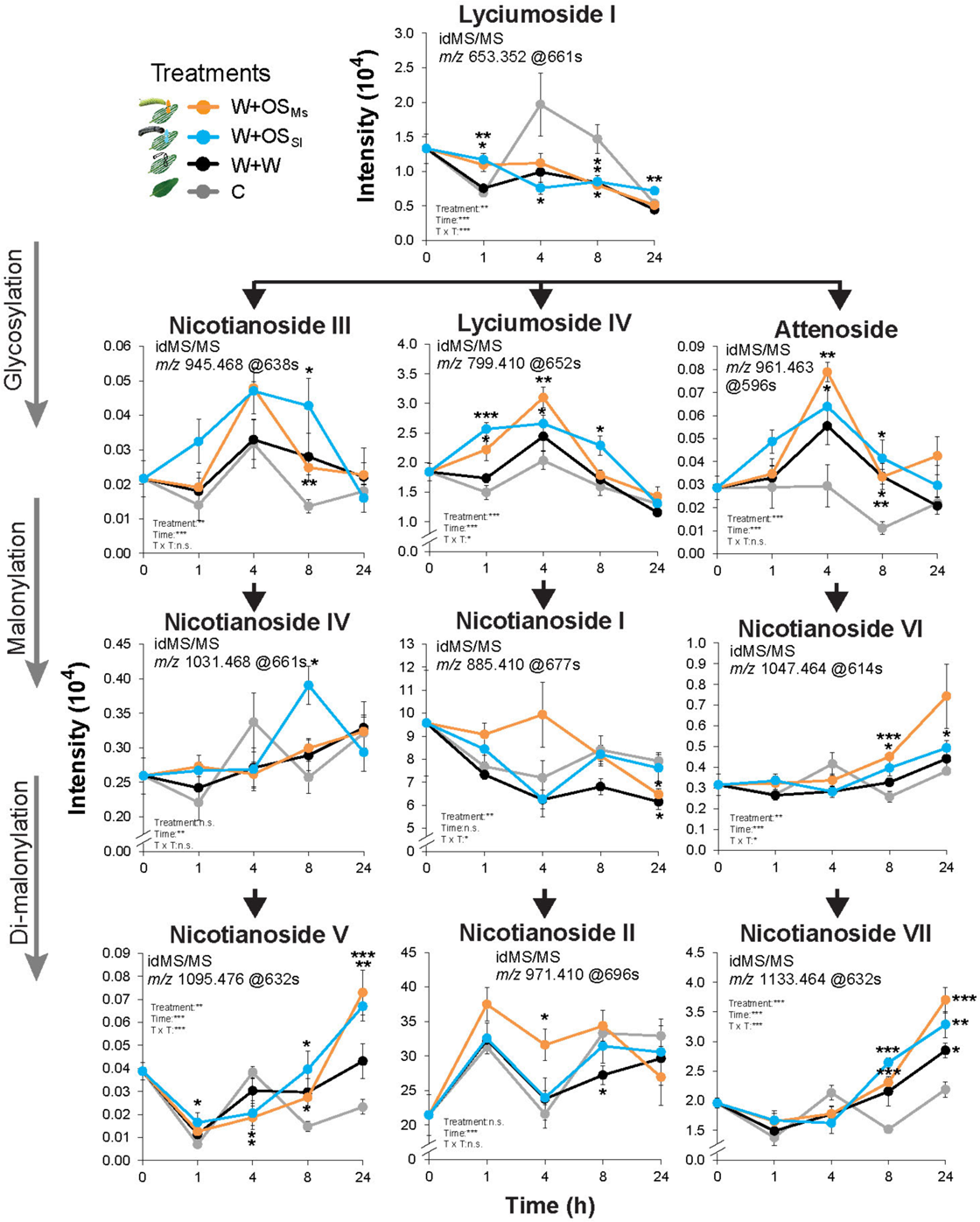
Accumulation of 17-HGL-DTGs in *N. attenuata* leaves in response to simulated herbivory. A pathway scale illustration of induction of 17-HGL-DTGs by simulated herbivory (W+OS: see legend) in which MS/MSs corresponding to most abundant HGL-DTGs were presented. Asterisks indicate significant differences between treatments (Two-way ANOVA followed by *post hoc* multiple comparisons, *P < 0.05, **P < 0.01, ***P < 0.001).

**Figure S6.**
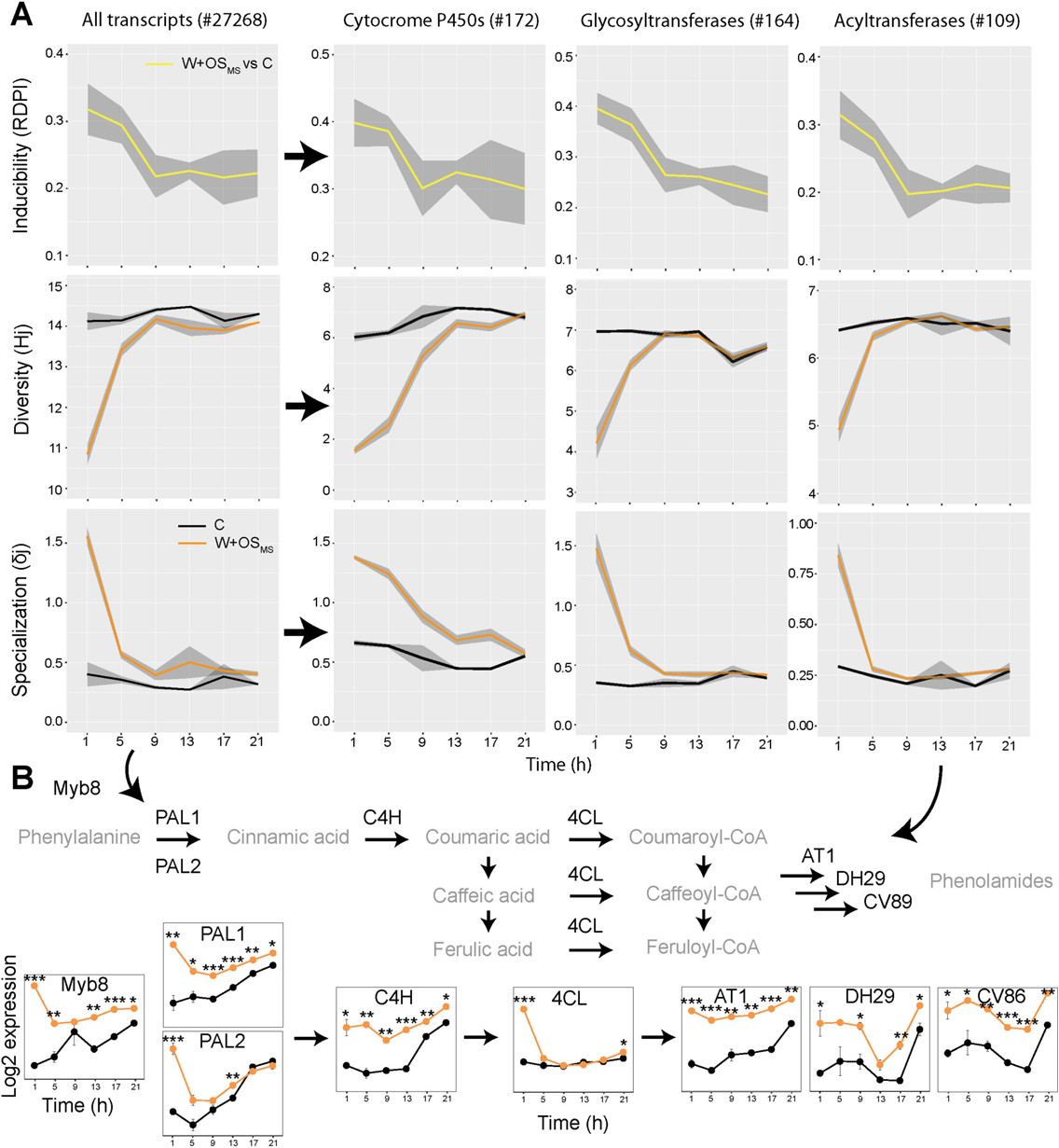
Information theory-based analysis of leaf transcriptome reconfigurations elicited by simulated *M. sexta* herbivory. (**A**) Transcriptome inducibility (RDPI), diversity (Hj) and specialization (δj) were calculated and visualized separately, where, RDPI was calculated by comparing Ms elicited leaves with undamaged controls (Microarray data were originally published in (*49*) and deposited in the National Center for Biotechnology Information Gene Expression Omnibus database (accession no. GSE30287)), Hj, transcriptome diversity was calculated by Shannon entropy of frequency distribution of each transcript and δj, transcriptome specialization was measured by the average specificity of each transcript by taking into consideration its frequency among samples. Subsets of genes corresponding to annotated metabolic gene families were further extracted and quantified by the three information theory descriptors. Ribbons refer to 95% confidence intervals. (**B**) Expression kinetics of key genes involved in the phenolamide biosynthesis. Asterisks indicate significant differences compared to controls (Two-way ANOVA followed by *post hoc* multiple comparisons, *P < 0.05, **P < 0.01, ***P < 0.001).

**Figure S7.**
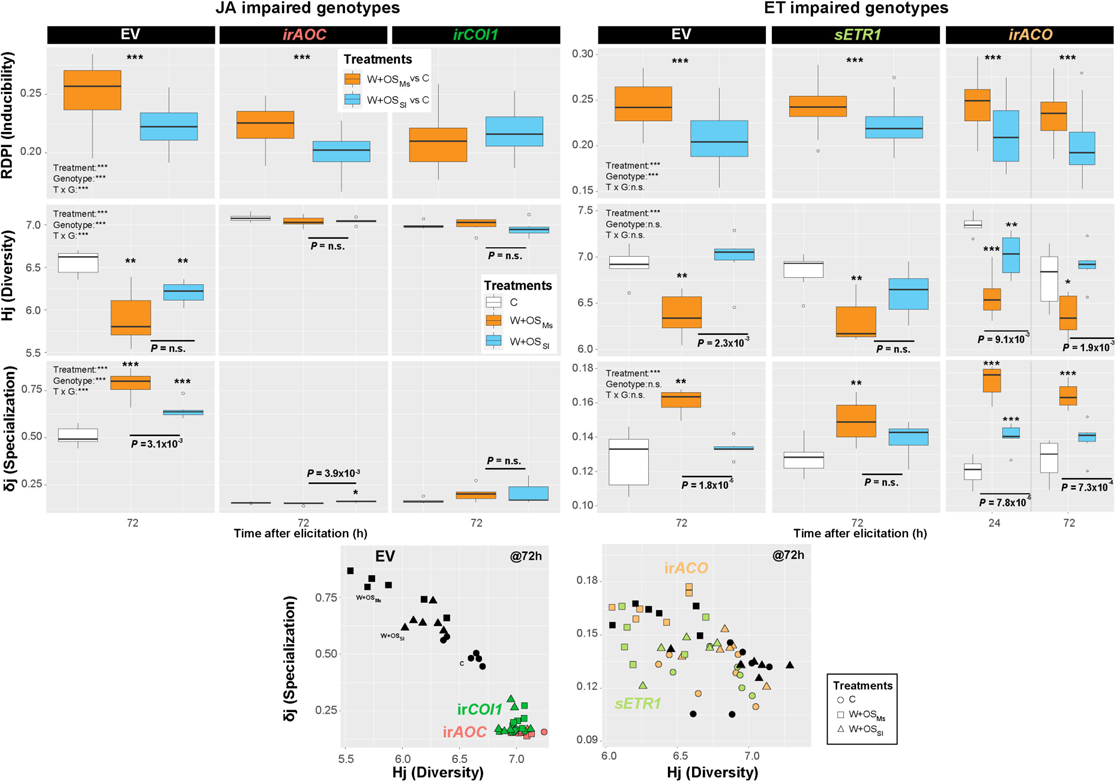
Use of transgenic plants reveals differential contribution of JA and ET signaling on herbivore species-specific modulations of specificity and diversity of leaf secondary metabolite profiles. Information theory analysis of 697 MS/MSs in jasmonate biosynthesis and perception-impaired lines (ir*AOC* and ir*COI1,* respectively) and 585 MS/MSs in the ethylene biosynthesis and perception-impaired lines (ir*ACO* and *sETR1,* respectively) as well as empty vector (EV) control plants elicited by the two simulated herbivory treatments (W+OS, see legend), demonstrate that jasmonates function as key determinants of metabolic plasticity whereas ethylene signaling modifies these herbivore-specific responses. Asterisks indicate significant differences between treatments (Two-way ANOVA followed by *post hoc* multiple comparisons, *P < 0.05, **P < 0.01, ***P < 0.001). Scatterplots of diversity against specialization in jasmonate- and ethylene-impaired transgenic lines at 72 h were additionally visualized in 2 dimensions. Colors indicate different transgenic lines and linear regression models were calculated for each genotype. Different symbols correspond to different treatments, triangle, W+OS_Sl_; rectangular, W+OS_Ms_; circle, C.

**Figure S8.**
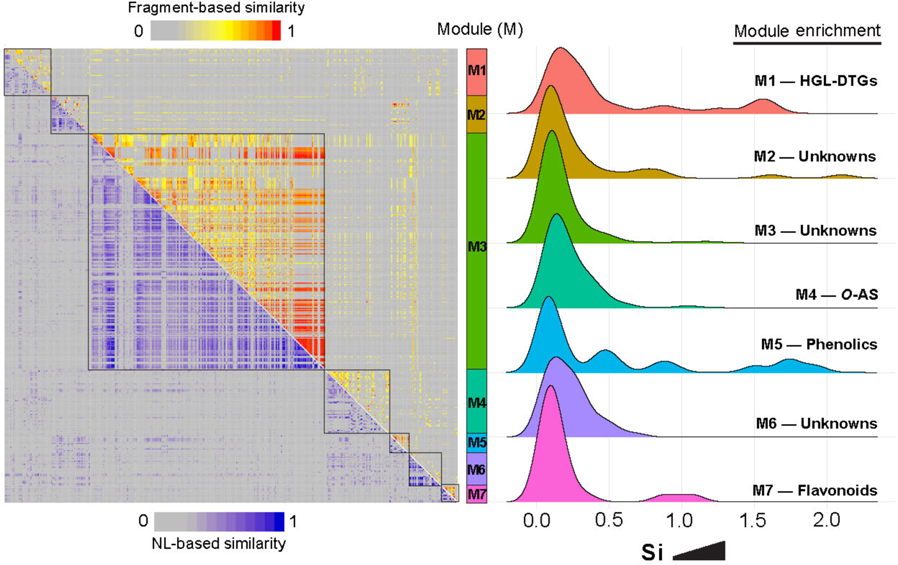
Metabolic classes extracted from the biclustering classification of MS/MSs contribute differentially to herbivore species-specific responses in JA-impaired transgenic plants. Biclustering analysis was used to classify MS/MSs according to structural similarities using two structural similarity scores, one based on shared fragments among spectra (NDP similarity), whereas the other scored shared common NLs among spectra (NL similarity). The biclustering analysis resulted in 7 modules depicted in different colors. Density plot of metabolic specificity (Si) of the whole dataset are depicted next to each module in which module enrichment is shown. A leptokurtic Si distribution of a module indicates certain metabolites with high metabolic specificity among plant genotypes or in response to different herbivores are grouped in this module. The Si distribution highlights metabolic groups contributing to high metabolic specificities that require JA signaling, such as HGL-DTGs, phenolics, etc.

**Figure S9.**
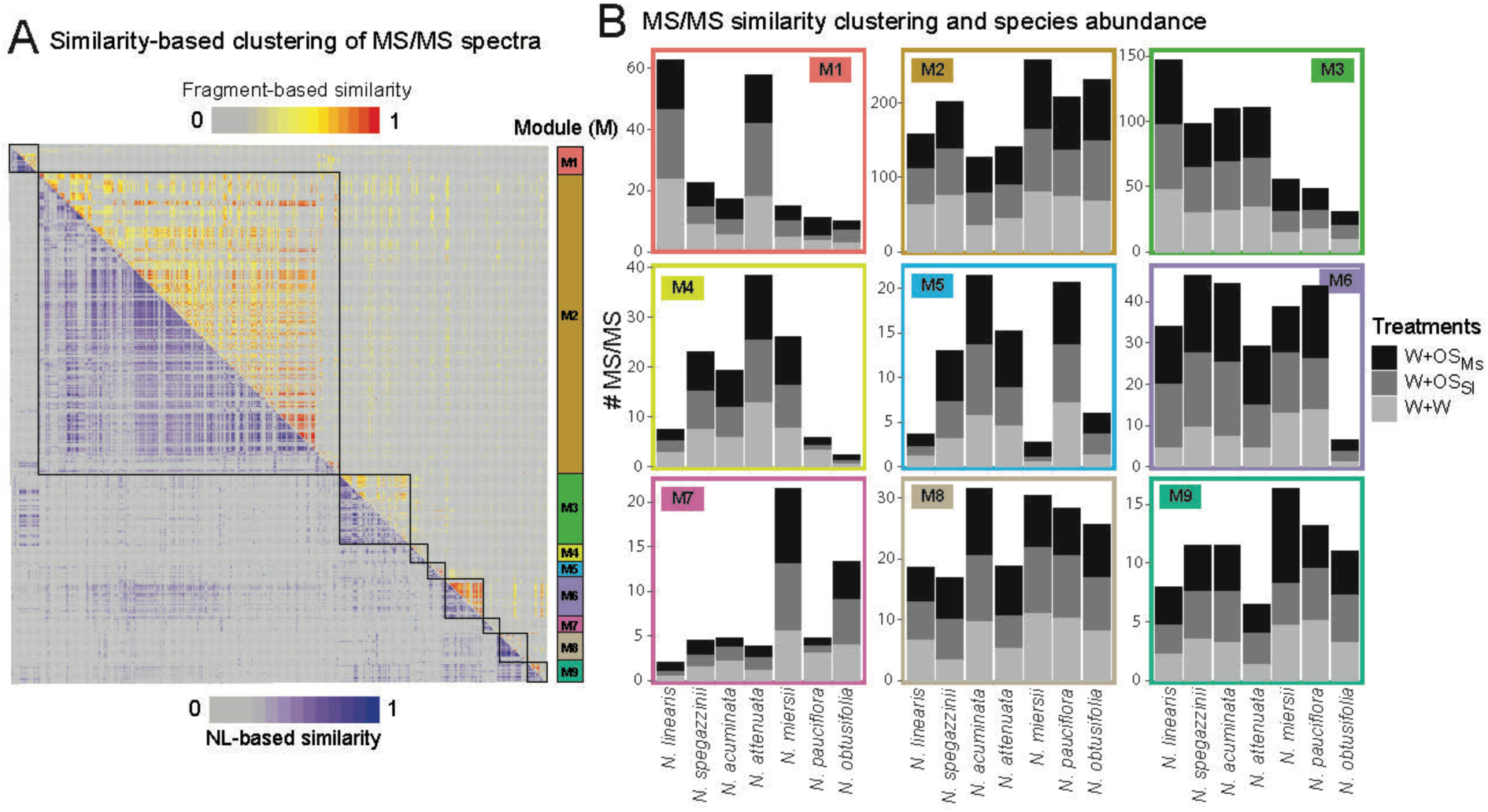
Metabolic classes extracted from the biclustering classification of MS/MSs differ in their herbivore species-specific reconfigurations of metabolic compositions among 7 closely related *Nicotiana* species. (A) Biclustering analysis to classify 939 MS/MSs in closely-related *Nicotiana* species. The biclustering analysis resulted in 9 modules depicted in different colors. **(B)** Relative contribution of each treatment to the MS/MSs associated with a given module among 7 species. Different treatments are shown as different colors.

**Figure S10.**
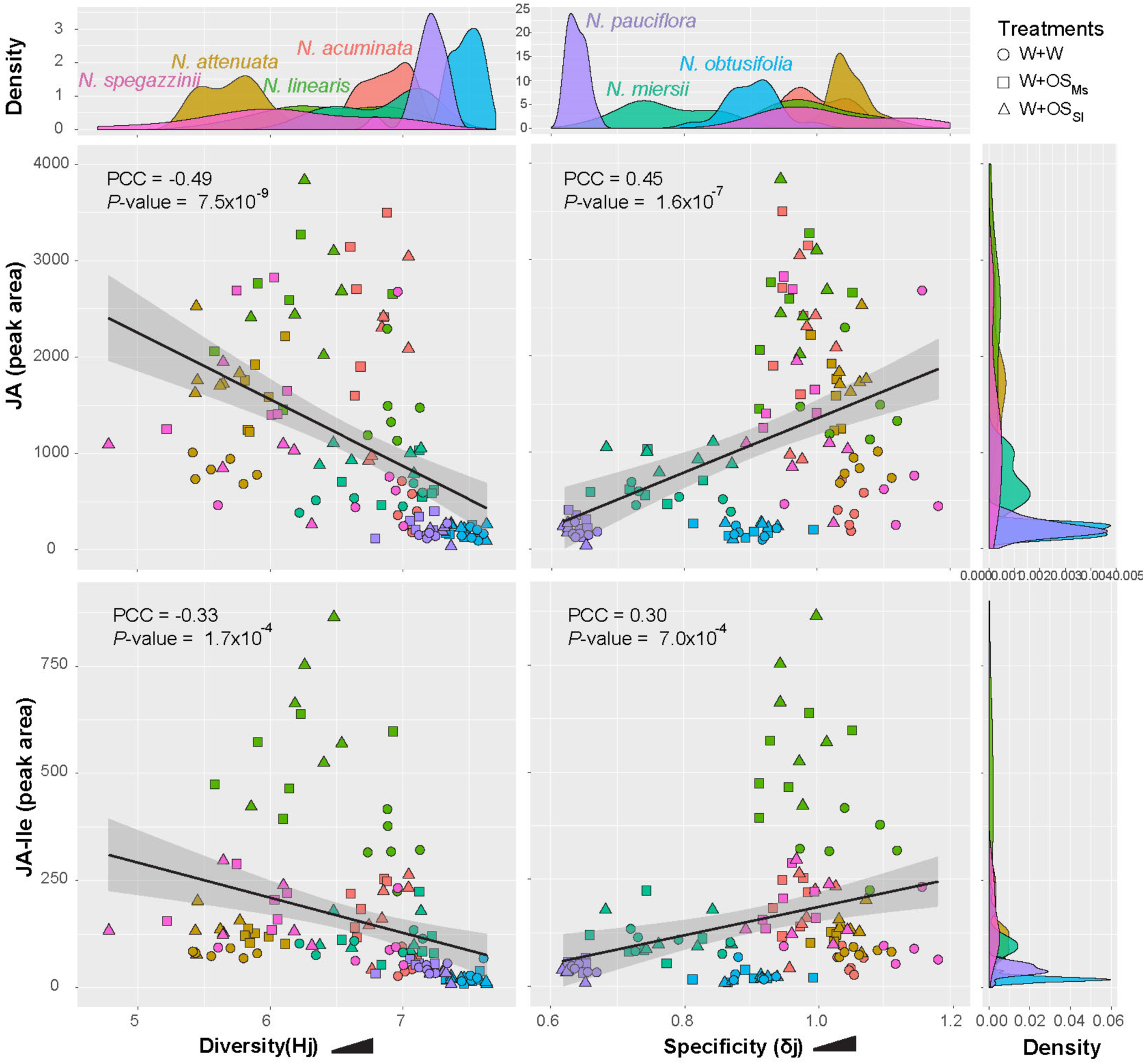
Amplitude of OS-specific induced JA and JA-Ile bursts correlates positively with the increase metabolic profile specialization across closely-related *Nicotiana* species. Species-level diversity (Hj) is negatively correlated with JA and JA-Ile, whereas specialization (δj) is positively correlated with JA and JA-Ile. A linear regression model is shown for each correlation. Different colors indicate different species and symbols correspond to different treatments. Density plots for each species are shown next to the scatter plots. PCC, pearson correlation coefficient.

**Figure S11.**
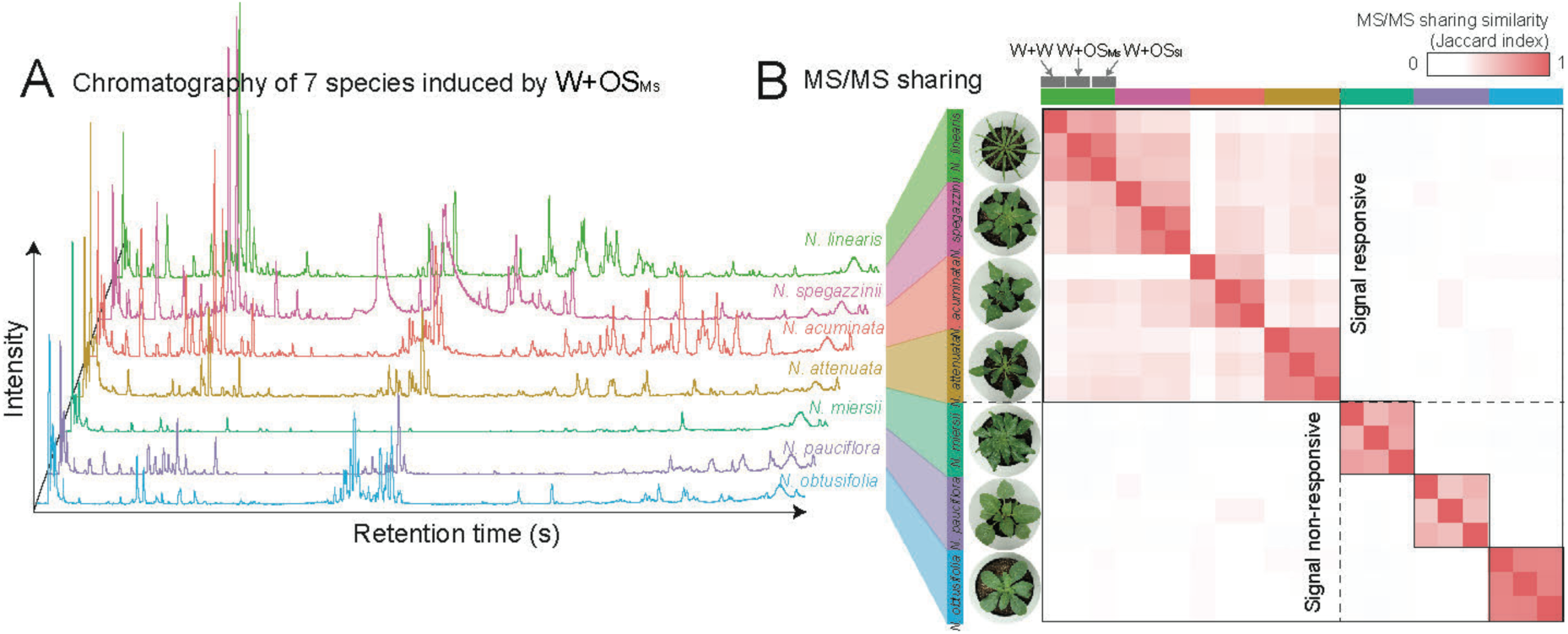
MS/MS sharing analysis reveals herbivory-induced metabolic interdependencies among *Nicotiana* species. (A) Representative chromatography of 7 *Nicotiana* species that were subjected to W+OS_Ms_ treatment. Different colors indicate different plant species. **(B)** Heat map matrix visualizing MS/MS sharing among 7 *Nicotiana* species subjected to W+W, W+ OS_Ms_ and W+ OS_Sl_ treatments as measured using the Jaccard index. Pictures of different species at rosette stage are shown next to the heat map.

**Figure S12.**
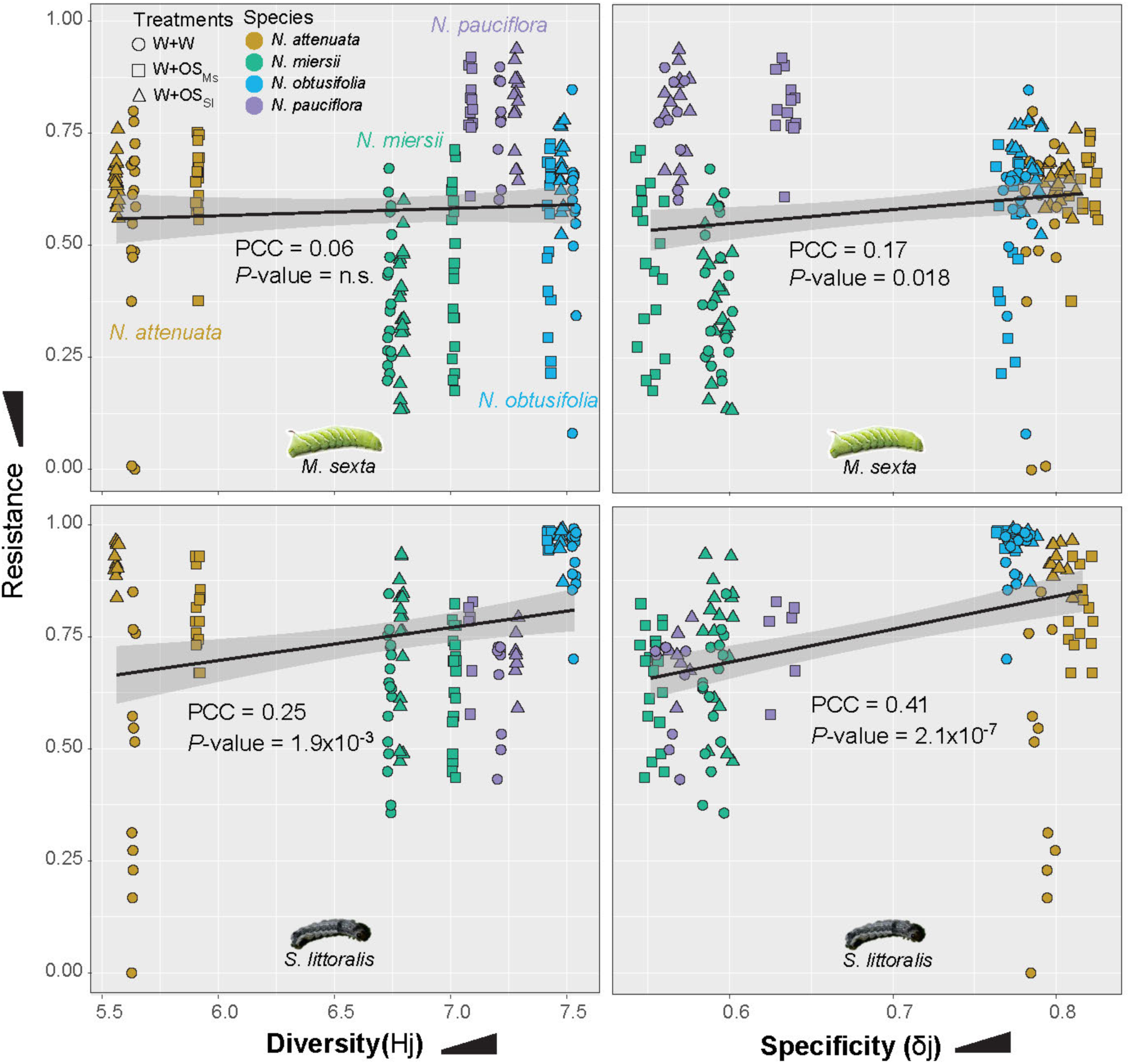
Herbivory-induced increases in the specialization of leaf metabolic profiles correlate positively with resistance to the generalist *S. littoralis* and the specialist *M. sexta* herbivores. Four of the *Nicotiana* species were selected for test of resistance to *M. sexta* (upper panels) and *S. littoralis* (lower panels) as previously described (*54*). For each plant species, rosette stage leaves were wounded and treated with either OS_Ms_, OS_Sl_, or water. 24 h after elicitation, leaves which had been differentially reconfigured in their metabolomes by different treatments were detached for herbivore feeding assays. Induced metabolome diversity (Hj) and specialization (δj) were first averaged and then mapped with linear regression models against a plant’s resistance to *M. sexta* and *S. littoralis* attack, respectively. Correlational analysis of metabolic diversity and specialization with herbivory resistance is shown adjacent to the regression line. Different colors indicate different species and symbols correspond to different treatments.

**Figure S13.**
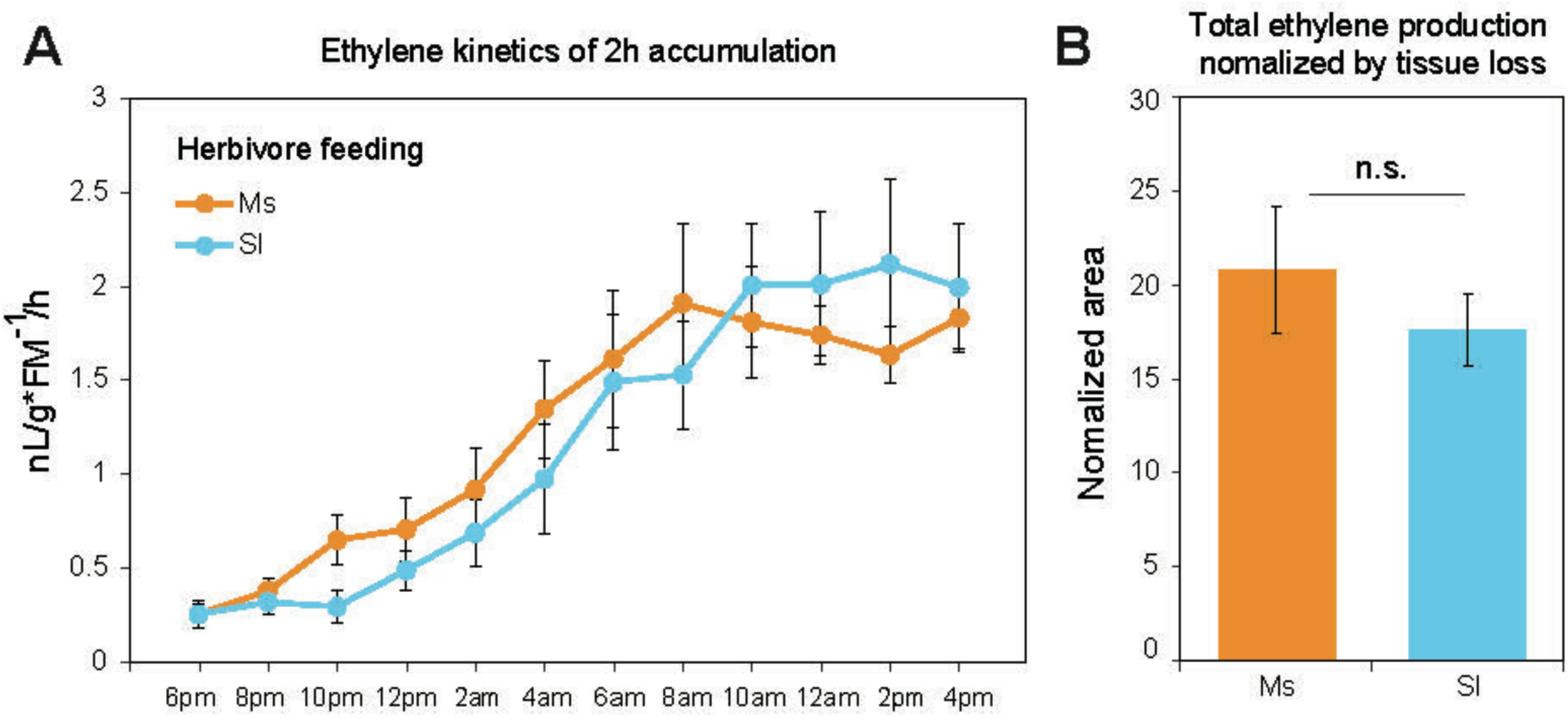
Real-time ethylene emission elicited by continuous herbivore feeding. (**A**) Leaves of *N. attenuata* were detached and fed on by *M. sexta* or *S. littoralis* larvae. Ethylene accumulations for each 2-h time interval were quantified. (**B**) Total ethylene produced after 24 h of continuous herbivore feeding was normalized to the tissue mass consumed by herbivores.

**Table S1.** List of the 52 neutral losses used for neutral loss similarity calculation

| No. | Loss name | Loss formula | Mass difference ± |
| --- | --- | --- | --- |
| 1 | Alkane chains, waxes, fatty acids, methylation | CH<sub>2</sub> | 14.0157 |
| 2 | Methane | CH<sub>4</sub> | 16.0313 |
| 3 | Ammonium adduct/neutral ammonium loss | NH<sub>3</sub> | 17.0265 |
| 4 | Water addition/loss | H<sub>2</sub>O | 18.0106 |
| 5 | Salt adduct | K<sup>+</sup> to NH<sub>4</sub><sup>+</sup> | 20.9293 |
| 6 | Salt adduct | Na<sup>+</sup> to H<sup>+</sup> | 21.9819 |
| 7 | Ethine | C<sub>2</sub>H<sub>2</sub> | 26.0157 |
| 8 | Carbon monoxide | CO | 27.9949 |
| 9 | Ethene | C<sub>2</sub>H<sub>4</sub> | 28.0313 |
| 10 | Methylimine | CH<sub>3</sub>N | 29.0266 |
| 11 | Formaldehyde | CH<sub>2</sub>O | 30.0106 |
| 12 | Methylamine | CH<sub>5</sub>N | 31.0422 |
| 13 | Sulfur | S | 31.9721 |
| 14 | Hydrogen sulfide | H<sub>2</sub>S | 33.9877 |
| 15 | Salt adduct | K<sup>+</sup> to H<sup>+</sup> | 37.9559 |
| 16 | Ketene | C<sub>2</sub>H<sub>2</sub>O | 42.0106 |
| 17 | Propylation | C<sub>3</sub>H<sub>6</sub> | 42.0470 |
| 18 | Cyanic acid | CHNO | 43.0058 |
| 19 | Carbon dioxide | CO<sub>2</sub> | 43.9898 |
| 20 | Formic acid | CH<sub>2</sub>O<sub>2</sub> | 46.0055 |
| 21 | Butylation | C<sub>4</sub>H<sub>8</sub> | 56.0626 |
| 22 | Trimethylamine | C<sub>3</sub>H<sub>9</sub>N | 59.0735 |
| 23 | Acetic acid | $C_2H_4O_2$ | 60.0211 |
| 24 | Urea | $CH_4N_2O$ | 60.0324 |
| 25 | Sulfur dioxide | $SO_2$ | 63.9619 |
| 26 | Pentene | $C_5H_8$ | 68.0626 |
| 27 | Propionic acid | $C_3H_6O_2$ | 74.0368 |
| 28 | Benzene | $C_6H_6$ | 78.0470 |
| 29 | Sulfur trioxide | $SO_3$ | 79.9568 |
| 30 | Malonic anhydride | $C_3H_2O_3$ | 86.0004 |
| 31 | Isobutyric acid | $C_4H_8O_2$ | 88.0517 |
| 32 | Putrescine | $C_4H_{12}N_2$ | 88.1000 |
| 33 | Sulfuric acid | $H_2SO_4$ | 97.9674 |
| 34 | Phosphoric acid | $H_3PO_4$ | 97.9769 |
| 35 | Methyl butanoic acid | $C_5H_{10}O_2$ | 102.0618 |
| 36 | Malonic acid | $C_3H_4O_4$ | 104.0110 |
| 37 | Methyl pentanoic acid | $C_6H_{12}O_2$ | 116.0861 |
| 38 | Dihydrogen vinyl phosphate | $C_2H_5O_4P$ | 123.9926 |
| 39 | Pentose equivalent | $C_5H_8O_4$ | 132.0423 |
| 40 | Spermidine | $C_7H_{19}N_3$ | 145.1579 |
| 41 | Deoxy-Hexose equivalent | $C_6H_{10}O_4$ | 146.0579 |
| 42 | Hexose-$H_2O$ | $C_6H_{10}O_5$ | 162.0528 |
| 43 | Deoxy-Hexose equivalent | $C_6H_{12}O_5$ | 164.0685 |
| 44 | Glucuronic acid-$H_2O$ | $C_6H_8O_6$ | 176.0321 |
| 45 | Hexose | $C_6H_{12}O_6$ | 180.0634 |
| 46 | Glucuronic acid | $C_6H_{10}O_7$ | 194.0427 |
| 47 | Acyl-fructose | $C_8H_{12}O_6$ | 204.0655 |
| 48 | Sinapic acid - H<sub>2</sub>O | C<sub>11</sub>H<sub>10</sub>O<sub>4</sub> | 206.0579 |
| 49 | Glutathione+O-H<sub>2</sub>O | C<sub>10</sub>H<sub>15</sub>N<sub>3</sub>O<sub>6</sub>S | 305.0682 |
| 50 | Glutathione | C<sub>10</sub>H<sub>17</sub>N<sub>3</sub>O<sub>6</sub>S | 307.0838 |
| 51 | Sucrose-H<sub>2</sub>O | C<sub>12</sub>H<sub>20</sub>O<sub>10</sub> | 324.1057 |
| 52 | Sucrose | C<sub>12</sub>H<sub>22</sub>O<sub>11</sub> | 342.1162 |

## References and Notes

1. A. Kessler, A. Kalske, Plant secondary metabolite diversity and species interactions. Annual Review of Ecology, Evolution, and Systematics 49, 115 (2018).

2. L. A. Dyer et al., Modern approaches to study plant-insect interactions in chemical ecology. Nat Rev Chem 2, 50 (Jun, 2018).

3. P. W. Sherman, The levels of analysis. Anim Behav 36, 616 (Apr, 1988).

4. M. C. Schuman, I. T. Baldwin, The layers of plant responses to insect herbivores. Annual Review of Entomology 61, 373 (2016).

5. B. E. Sedio, Recent breakthroughs in metabolomics promise to reveal the cryptic chemical traits that mediate plant community composition, character evolution and lineage diversification. New Phytol. 214, 952 (May, 2017).

6. C. E. Shannon, A mathematical theory of communication. Bell Syst Tech J 27, 379 (1948).

7. R. E. Ulanowicz, Information theory in ecology. Computers & chemistry 25, 393 (Jul, 2001).

8. A. Borst, F. E. Theunissen, Information theory and neural coding. Nature neuroscience 2, 947 (Nov, 1999).

9. T. D. Schneider, D. N. Mastronarde, Fast multiple alignment of ungapped DNA sequences using information theory and a relaxation method. Discrete Appl Math 71, 259 (Dec 5, 1996).

10. O. Martinez, M. H. Reyes-Valdes, Defining diversity, specialization, and gene specificity in transcriptomes through information theory. Proceedings of the National Academy of Sciences of the United States of America 105, 9709 (Jul 15, 2008).

11. D. Li, S. Heiling, I. T. Baldwin, E. Gaquerel, Illuminating a plant’s tissue-specific metabolic diversity using computational metabolomics and information theory. Proceedings of the National Academy of Sciences of the United States of America 113, E7610 (Nov 22, 2016).

12. D. Mckey, Adaptive patterns in alkaloid physiology. Am Nat 108, 305 (1974).

13. D. Mckey, *The distribution of secondary compounds within plants*. (Academic, New York, 1979), pp. 55–133.

14. D. F. Rhoades, *Evolution of plant chemical defense against herbivores*. (Academic, New York, 1979), pp. 1–55.

15. F. R. Adler, R. Karban, Defended fortresses or moving targets? Another model of inducible defenses inspired by military metaphors. Am Nat 144, 813 (Nov, 1994).

16. P. Feeny, Plant apparency and chemical defense. Recent Advances in Phytochemistry (Springer, Boston, MA, Biochemical Interaction Between Plants and Insects, 1976), vol. 10, pp. 1–40.

17. J. P. Bryant, F. S. Chapin, D. R. Klein, Carbon nutrient balance of boreal plants in relation to vertebrate herbivory. Oikos 40, 357 (1983).

18. P. D. Coley, J. P. Bryant, F. S. Chapin, Resource availability and plant antiherbivore defense. Science 230, 895 (1985).

19. W. E. Loomis, Growth-differentiation balance vs carbohydrate-nitrogen ratio. Proceedings of the American Society for Horticultural Science 29, 240 (1932).

20. R. A. Keith, T. Mitchell-Olds, Testing the optimal defense hypothesis in nature: Variation for glucosinolate profiles within plants. PloS one 12, (Jul 21, 2017).

21. A. L. Godschalx, L. Stady, B. Watzig, D. J. Ballhorn, Is protection against florivory consistent with the optimal defense hypothesis? BMC plant biology 16, (Jan 28, 2016).

22. A. C. McCall, J. A. Fordyce, Can optimal defence theory be used to predict the distribution of plant chemical defences? J Ecol 98, 985 (Sep, 2010).

23. J. N. Holland, S. A. Chamberlain, K. C. Horn, Optimal defence theory predicts investment in extrafloral nectar resources in an ant-plant mutualism. J Ecol 97, 89 (Jan, 2009).

24. V. Radhika, C. Kost, S. Bartram, M. Heil, W. Boland, Testing the optimal defence hypothesis for two indirect defences: extrafloral nectar and volatile organic compounds. Planta 228, 449 (Aug, 2008).

25. T. E. Ohnmeiss, I. T. Baldwin, Optimal Defense theory predicts the ontogeny of an induced nicotine defense. Ecology 81, 1765 (Jul, 2000).

26. E. K. Barto, D. Cipollini, Testing the optimal defense theory and the growth-differentiation balance hypothesis in *Arabidopsis thaliana*. Oecologia 146, 169 (Dec, 2005).

27. M. Wink, Evolution of secondary metabolites from an ecological and molecular phylogenetic perspective. Phytochemistry 64, 3 (Sep, 2003).

28. B. D. Moore, R. L. Andrew, C. Kulheim, W. J. Foley, Explaining intraspecific diversity in plant secondary metabolites in an ecological context. New Phytol. 201, 733 (Feb, 2014).

29. L. A. Richards et al., Phytochemical diversity drives plant-insect community diversity. Proceedings of the National Academy of Sciences of the United States of America 112, 10973 (Sep 1, 2015).

30. G. S. Fraenkel, The raison d’etre of secondary plant substances. Science 129, 1466 (May 29, 1959).

31. A. A. Agrawal, A. P. Hastings, M. T. Johnson, J. L. Maron, J. P. Salminen, Insect herbivores drive real-time ecological and evolutionary change in plant populations. Science 338, 113 (Oct 5, 2012).

32. A. Kempel, M. Schadler, T. Chrobock, M. Fischer, M. van Kleunen, Tradeoffs associated with constitutive and induced plant resistance against herbivory. Proceedings of the National Academy of Sciences of the United States of America 108, 5685 (Apr 5, 2011).

33. A. A. Agrawal, P. M. Gorski, D. W. Tallamy, Polymorphism in plant defense against herbivory: Constitutive and induced resistance in *Cucumis sativus*. J. Chem. Ecol. 25, 2285 (Oct, 1999).

34. I. T. Baldwin, Jasmonate-induced responses are costly but benefit plants under attack in native populations. Proceedings of the National Academy of Sciences of the United States of America 95, 8113 (Jul 7, 1998).

35. S. Y. Strauss, J. A. Rudgers, J. A. Lau, R. E. Irwin, Direct and ecological costs of resistance to herbivory. Trends in ecology & evolution 17, 278 (Jun, 2002).

36. D. Li, I. T. Baldwin, E. Gaquerel, Navigating natural variation in herbivory-induced secondary metabolism in coyote tobacco populations using MS/MS structural analysis. Proceedings of the National Academy of Sciences of the United States of America 112, E4147 (Jul 28, 2015).

37. E. Gaquerel, M. Stitz, M. Kallenbach, I. T. Baldwin, Jasmonate signaling in the field, part II: insect-guided characterization of genetic variations in jasmonate-dependent defenses of transgenic and natural *Nicotiana attenuata* populations. Methods Mol Biol 1011, 97 (2013).

38. P. B. Pelser et al., Frequent gain and loss of pyrrolizidine alkaloids in the evolution of *Senecio* section *Jacobaea* (Asteraceae). Phytochemistry 66, 1285 (Jun, 2005).

39. T. Züst et al., Natural enemies drive geographic variation in plant defenses. Science 338, 116 (Oct 5, 2012).

40. F. Valladares, D. Sanchez-Gomez, M. A. Zavala, Quantitative estimation of phenotypic plasticity: bridging the gap between the evolutionary concept and its ecological applications. J Ecol 94, 1103 (Nov, 2006).

41. C. Hettenhausen, I. T. Baldwin, J. Q. Wu, *Nicotiana attenuata* MPK4 suppresses a novel jasmonic acid (JA) signaling-independent defense pathway against the specialist insect *Manduca sexta*, but is not required for the resistance to the generalist *Spodoptera littoralis*. New Phytol. 199, 787 (Aug, 2013).

42. H. Kaur, N. Heinzel, M. Schottner, I. T. Baldwin, I. Galis, R2R3-NaMYB8 regulates the accumulation of phenylpropanoid-polyamine vonjugates, which are essential for local and systemic defense against insect herbivores in *Nicotiana attenuata*. Plant physiology 152, 1731 (Mar, 2010).

43. N. Onkokesung et al., MYB8 controls inducible phenolamide levels by activating three novel hydroxycinnamoyl-coenzyme A:polyamine transferases in *Nicotiana attenuata*. Plant physiology 158, 389 (Jan, 2012).

44. S. Heiling et al., Jasmonate and ppHsystemin regulate key malonylation steps in the biosynthesis of 17-hydroxygeranyllinalool diterpene glycosides, an abundant and effective direct defense against herbivores in *Nicotiana attenuata*. The Plant cell 22, 273 (Jan, 2010).

45. E. S. McCloud, I. T. Baldwin, Herbivory and caterpillar regurgitants amplify the wound-induced increases in jasmonic acid but not nicotine in *Nicotiana sylvestris*. Planta 203, 430 (Dec, 1997).

46. E. Gaquerel, S. Heiling, M. Schoettner, G. Zurek, I. T. Baldwin, Development and validation of a liquid chromatography-electrospray ionization-time-of-flight mass spectrometry method for induced changes in *Nicotiana attenuata* leaves during simulated herbivory. J Agr Food Chem 58, 9418 (Sep 8, 2010).

47. J. Gulati, S. G. Kim, I. T. Baldwin, E. Gaquerel, Deciphering herbivory-induced gene-to-metabolite dynamics in *Nicotiana attenuata* tissues using a multifactorial approach. Plant physiology 162, 1042 (Jun, 2013).

48. H. Takahashi et al., Dynamics of time-lagged gene-to-metabolite networks of *Escherichia coli* elucidated by integrative omics approach. Omics 15, 15 (Jan, 2011).

49. S. G. Kim, F. Yon, E. Gaquerel, J. Gulati, I. T. Baldwin, Tissue specific diurnal rhythms of metabolites and their regulation during herbivore attack in a native tobacco, *Nicotiana attenuata*. PloS one 6, (Oct 18, 2011).

50. M. Kallenbach, G. Bonaventure, P. A. Gilardoni, A. Wissgott, I. T. Baldwin, *Empoasca* leafhoppers attack wild tobacco plants in a jasmonate-dependent manner and identify jasmonate mutants in natural populations. Proceedings of the National Academy of Sciences of the United States of America 109, E1548 (Jun 12, 2012).

51. T. H. Goodspeed, The genus Nicotiana. (Chronica Botanica Company, Waltham, Mass., 1954), vol. 16, pp. 102–35.

52. J. J. Clarkson, L. J. Kelly, A. R. Leitch, S. Knapp, M. W. Chase, Nuclear glutamine synthetase evolution in *Nicotiana*: Phylogenetics and the origins of allotetraploid and homoploid (diploid) hybrids. Mol Phylogenet Evol 55, 99 (Apr, 2010).

53. W. W. Zhou et al., Evolution of herbivore-induced early defense signaling was shaped by genome wide duplications in *Nicotiana*. eLife 5, (Nov 4, 2016).

54. S. Q. Xu, W. W. Zhou, S. Pottinger, I. T. Baldwin, Herbivore associated elicitor-induced defences are highly specific among closely related *Nicotiana* species. BMC plant biology 15, (Jan 16, 2015).

55. J. R. Platt, Strong inference: certain systematic methods of scientific thinking may produce much more rapid progress than others. Science 146, 347 (1964).

56. N. Stamp, Out of the quagmire of plant defense hypotheses. Q Rev Biol 78, 23 (Mar, 2003).

57. U. Schittko, C. A. Preston, I. T. Baldwin, Eating the evidence? *Manduca sexta* larvae can not disrupt specific jasmonate induction in *Nicotiana attenuata* by rapid consumption. Planta 210, 343 (Jan, 2000).

58. H. J. Guo et al., A porin-like protein from oral secretions of *Spodoptera littoralis* larvae induces defense-related early events in plant leaves. Insect Biochem Molec 43, 849 (Sep, 2013).

59. L. Mack, P. Gros, J. Burkhardt, K. Seifert, Elicitors of tansy volatiles from cotton leafworm larval oral secretion. Phytochemistry 96, 158 (Dec, 2013).

60. R. Sutter, C. Muller, Mining for treatment-specific and general changes in target compounds and metabolic fingerprints in response to herbivory and phytohormones in *Plantago lanceolata*. New Phytol. 191, 1069 (2011).

61. R. Schweiger, A. M. Heise, M. Persicke, C. Muller, Interactions between the jasmonic and salicylic acid pathway modulate the plant metabolome and affect herbivores of different feeding types. Plant Cell and Environment 37, 1574 (Jul, 2014).

62. C. C. von Dahl et al., Tuning the herbivore-induced ethylene burst: the role of transcript accumulation and ethylene perception in *Nicotiana attenuata*. Plant Journal 51, 293 (Jul, 2007).

63. R. A. Winz, I. T. Baldwin, Molecular interactions between the specialist herbivore *Manduca sexta* (Lepidoptera, Sphingidae) and its natural host Nicotiana attenuata. IV. Insect-induced ethylene reduces jasmonate-induced nicotine accumulation by regulating putrescine N-methyltransferase transcripts. Plant physiology 125, 2189 (Apr, 2001).

64. M. P. Ayres, T. P. Clausen, S. F. MacLean, A. M. Redman, P. B. Reichardt, Diversity of structure and antiherbivore activity in condensed tannins. Ecology 78, 1696 (Sep, 1997).

65. R. H. Whittaker, Evolution and measurement of species diversity. Taxon 21, 213 (1972).

66. P. R. Ehrlich, P. H. Raven, Butterflies and plants - a study in coevolution. Evolution 18, 586 (1964).

67. A. A. Agrawal et al., Evidence for adaptive radiation from a phylogenetic study of plant defenses. Proceedings of the National Academy of Sciences of the United States of America 106, 18067 (Oct 27, 2009).

68. R. D. Firn, C. G. Jones, Natural products - a simple model to explain chemical diversity. Nat Prod Rep 20, 382 (Aug, 2003).

69. M. Schäfer, C. Brütting, I. T. Baldwin, M. Kallenbach, High-throughput quantification of more than 100 primary- and secondary-metabolites, and phytohormones by a single solid-phase extraction based sample preparation with analysis by UHPLC-HESI-MS/MS. Plant Methods 12, (May 26, 2016).

70. C. C. von Dahl, I. T. Baldwin, Deciphering the role of ethylene in plant-herbivore interactions. J Plant Growth Regul 26, 201 (Jun, 2007).

71. M. Pertea, D. Kim, G. M. Pertea, J. T. Leek, S. L. Salzberg, Transcript-level expression analysis of RNA-seq experiments with HISAT, StringTie and Ballgown. Nat Protoc 11, 1650 (Sep, 2016).

72. H. Horai et al., MassBank: a public repository for sharing mass spectral data for life sciences. Journal of mass spectrometry: JMS 45, 703 (Jul, 2010).

73. B. M. Tesson, R. Breitling, R. C. Jansen, DiffCoEx: a simple and sensitive method to find differentially coexpressed gene modules. BMC bioinformatics 11, (Oct 6, 2010).

